# Extrusion fountains are hallmarks of chromosome organization emerging upon zygotic genome activation

**DOI:** 10.1101/2023.07.15.549120

**Authors:** Aleksandra Galitsyna, Sergey V. Ulianov, Mariia Bazarevich, Nikolai S. Bykov, Marina Veil, Meijiang Gao, Kristina Perevoschikova, Mikhail S. Gelfand, Sergey V. Razin, Leonid Mirny, Daria Onichtchouk

## Abstract

The initiation of gene expression during development, known as zygotic genome activation (ZGA), is accompanied by massive changes in chromosome organization. However, the earliest events of chromosome folding and their functional roles remain unclear. Using Hi-C on zebrafish embryos, we discovered that chromosome folding begins early in development with the formation of "fountains", a novel element of chromosome organization. Emerging preferentially at enhancers, fountains exhibit an initial accumulation of cohesin, which later redistributes to CTCF sites at TAD borders. Knockouts of pioneer transcription factors driving ZGA enhancers result in the specific loss of fountains, establishing a causal link between enhancer activation and fountain formation. Polymer simulations demonstrate that fountains may arise as sites of facilitated cohesin loading, requiring two-sided but desynchronized loop extrusion, potentially caused by cohesin collisions with obstacles or internal switching. Moreover, we detected similar fountain patterns at enhancers in mouse cells. Fountains disappear upon acute cohesin depletion, as well as during mitosis, and reappear with cohesin loading in early G1. Altogether, fountains represent the first known enhancer-specific elements of chromosome organization and constitute starting points for chromosome folding during development, likely through facilitated cohesin loading.

## Introduction

Zygotic genome activation (ZGA) is a critical point in the development of multicellular organisms. It is largely driven by the binding of pioneer transcription factors that open up chromatin and activate enhancers, leading to the awakening of zygotic transcription (reviewed in ^1,2^). ZGA also coincides with dramatic reorganization of chromosome folding. In *Drosophila*, *Xenopus*, zebrafish, medaka, human, and mouse, the hallmarks of chromatin organization gradually emerge after ZGA, with largely featureless organization before ZGA ^3–10^. As observed by Hi-C in all vertebrates, these hallmarks include compartments at larger scales, as well as topologically associated domains (TADs), dots and stripes, at a sub-megabase scale. Central to the formation of these sub-megabase structures is the process of loop extrusion by cohesin. Interacting with extrusion barriers such as CTCF ^11,12^, cohesin forms highly dynamic ^13–15^ chromatin structures. Functionally, cohesin-mediated loop extrusion and its modulation by CTCF play key roles in interactions between distal enhancers and their target promoters, and are crucial for development and differentiation ^16–20^.

Although the establishment of specific chromosome organization during development coincides with the major ZGA wave, the causal relationship between these two processes remains unclear. In zebrafish, ZGA is driven by pioneering transcription factors (TFs) Pou5f3, Sox19b, and Nanog ^21,22^ (homologs of mouse pluripotency TFs ^23^), which open chromatin, establish early enhancers, promote H3K27 acetylation, and activate transcription ^24–26^.

The initial steps of chromosome folding may also be guided by pioneer TFs that can load cohesin or recruit insulators, such as CTCF. For example, in *Drosophila*, the removal of the pioneer factor Zelda, a zygotic genome activator, leads to the weakening and loss of a subset of domain boundaries ^3^. In zebrafish, cohesin and CTCF levels gradually increase after ZGA ^27^, but the factors and mechanisms responsible for their recruitment remain largely unknown. Overall, the mechanisms underlying this dramatic initial chromosome folding during development, and the roles of enhancers and pioneer TFs in this process, are still not fully understood.

Here, we generated Hi-C maps at several time points during early development for wild-type and mutant zebrafish embryos lacking various combinations of key pioneer transcription factors. Our analysis revealed that, upon ZGA, chromosomes begin to fold by forming "fountains," a new class of Hi-C features that we systematically detected and functionally characterized. Fountains emerging in early development resemble structures previously observed in Wapl-CTCF-depleted mouse cells ("plumes") ^28^, quiescent mouse thymocytes ("jets") ^29^, and in *C.elegans* ("fountains" ^30^, or "jets" ^31^, see discussion in Suppl. Information).

Surprisingly, fountains are enriched at early enhancers, rather than at active promoters or other functional genomic elements. Surrounded by genes active in early development, fountains transform into other elements of chromosome organization as development progresses. We hypothesized and demonstrated through simulations that these initial enhancer-specific structures are driven by facilitated cohesin loading at enhancers. TFs knockouts supported this mechanism, showing that disruption of enhancer activity leads to impaired fountain formation. Furthermore, we identified fountains as a collective signature of enhancers in mouse cells, suggesting that they represent a general enhancer-associated folding mechanism. Their cohesin-dependent nature is evident from their disappearance upon induced cohesin degradation, their absence during mitosis, and their gradual reemergence after mitosis. Consistently, fountain-like structures were also found to be cohesin-dependent in thymocytes and *C. elegans*. Overall, our findings indicate that chromosome folding during development is initiated by facilitated cohesin loading at enhancers, revealing a new role for enhancers as active participants in chromosome folding.

## Results

### Large-scale chromosome reorganization in early zebrafish embryogenesis

To characterize the dynamics of chromosome organization, we performed Hi-C for sperm and four stages of *Danio rerio* embryonic development: the last cell cycle before the major ZGA wave (2.75 hours post fertilization, hpf), late blastula (5.3 hpf, 2.3 hours after ZGA), 3-somite stage (11 hpf) and pharyngula (25 hpf) ^32^ (Fig. 1a).

**Figure 1.**
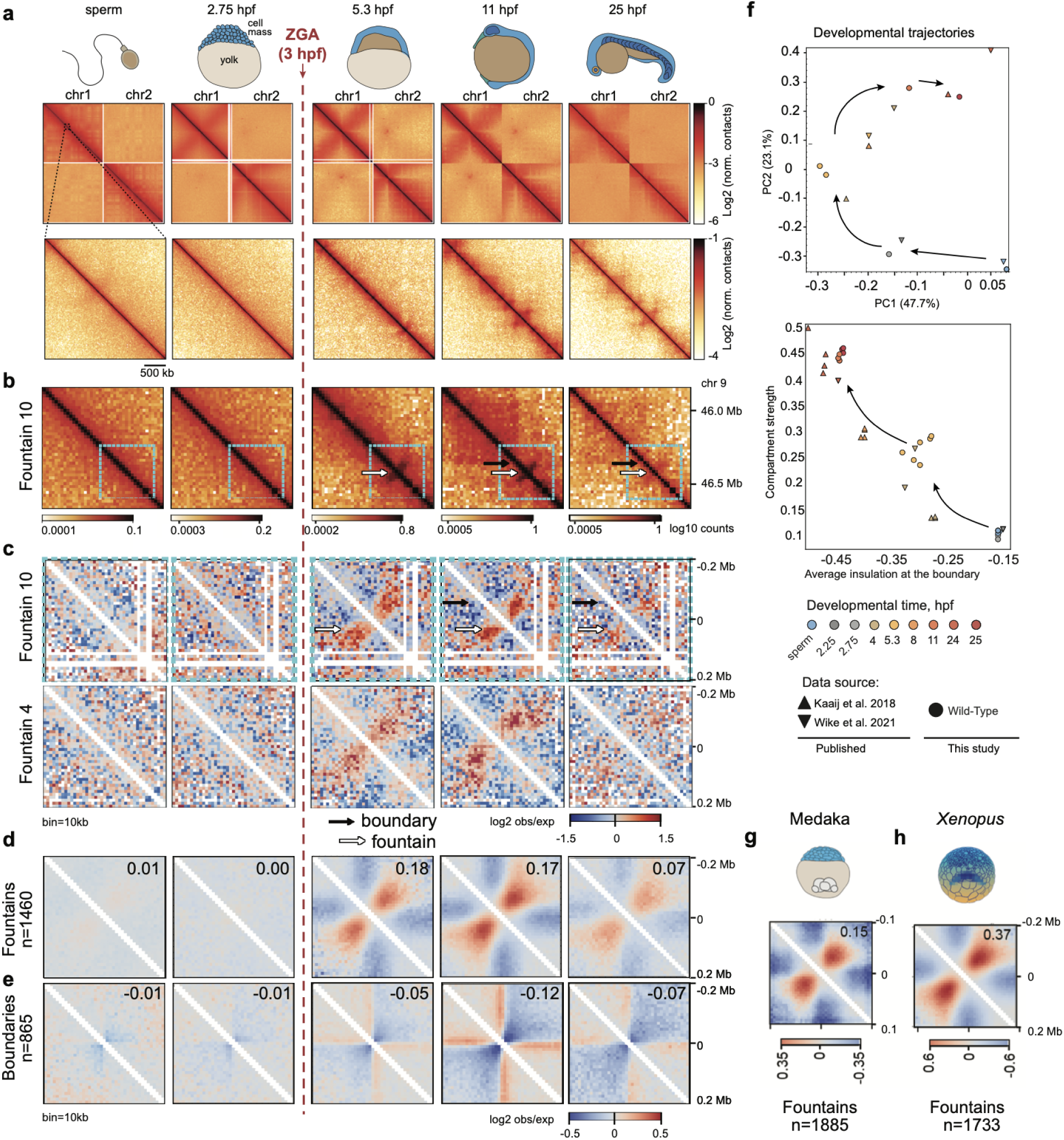
Structural features of chromatin of embryogenesis and emergence of fountains after ZGA. ZGA – zygotic genome activation; hpf – hours post fertilization; n – number of instances used for averaging; log2 norm. contacts – logarithm of Hi-C contacts normalized by iterative correction; log2 obs/exp – logarithm of observed Hi-C contacts divided by expected for each genomics separation; PC – principal component. Vertical dashed red line represents ZGA onset in zebrafish development. **a.** Representative examples of contact heatmaps showing dynamics of whole-chromosomal (top, chr1-2) and local contact patterns (bottom, chr10: 31,200,000-34,000,000). Dashed line symbolizes the scale of comparison between top and bottom but not the exact location. **b.** A representative example of a fountain (white arrow) on the contact heatmap after ZGA (5.3 hpf). A TAD boundary (black arrow) is detectable at 11 hpf. The region in the blue frame is shown in detail in (c, top). Bin size=20 kb. **c.** Examples of fountains in Hi-C maps normalized by expected. Top: the same region as in (b), bottom: another example (see Suppl. Data for more examples). Bin size=10 kb. Coordinates represent disances from the center of the Hi-C snippet. **d.** Average fountain (defined at 5.3 hpf, n=1460). The average fountain score is shown at the top right corner. Coordinates represent disance from the fountain base. **e.** Average TAD boundary (defined at 11 hpf, n=865). The average insulation score is shown at the top right corner. Coordinates represent disances from the boundary. **f.** Developmental trajectories based on stratum-adjusted correlation coefficient (SCC, *top*) and average insulation plotted against compartmental strength (*bottom*). Plots include datasets from three studies (^4,33^ and this work). Percentage numbers in the top plot label the variance explained by the PCA components. (**g, h**). Fountains at ZGA in other species. Fountains called using algorithm *fontanka* and reported in Suppl. Dataset 2. **g.** Average fountain observed in medaka fish based on 1855 fountains. Hi-C for developmental stage 10 (early blastula) from ^6^. **h.** Average fountain observed in *Xenopus tropicalis* based on 1733 fountains. Hi-C for developmental stage 11 (gastrula) from ^9^.

While Hi-C maps were largely featureless before ZGA, the hallmarks of chromosome organization started to emerge at 5.3 hpf. At the global level, we noticed the formation of compartments and a strong Rabl configuration, i.e., enrichment of interactions between centromeric regions (ED Fig. 1a,b). At the local level (<1Mb), after ZGA (5.3 hpf), we observed the formation of features, referred to below as “fountains” (Fig. 1b-d). Fountains are distinct from TADs, yet morph into TADs at much later stages (Fig. 1b-e). At 11 hpf, compartments and TADs became more pronounced, and practically fully formed as compared to 24 hpf stage (Fig. 1e, ED Fig. 1b,c). While early Hi-C studies of zebrafish development suggested presence of TADs and compartments before ZGA ^33^, later reports ^4^ as well as our data indicate absence of features in chromosome organization before ZGA (Fig. 1a-e), consistent with observations in other species ^3–10^. Fountains appear to be the earliest features emerging after ZGA, suggesting that chromosome folding starts by formation of fountains.

For comparison of our datasets with the published Hi-C data on *Danio rerio* development ^4,33^, we constructed “developmental trajectories” by two independent approaches, (i) pairwise correlation-based comparisons and embeddings of Hi-C maps into 2D space, and (ii) comparing datasets in their degrees of compartmentalization and insulation at TAD boundaries. Both approaches demonstrate that our datasets are well-aligned with the previous data. Interestingly, we observe that compartmentalization and insulation grow together, gradually and in synchrony with each other, during early development (Fig. 1f).

### Emergence of loop extrusion activity upon ZGA

Gradual increase in insulation (Fig. 1f, ED Fig. 1c) suggests the rising level of loop extrusion activity shortly after ZGA and its increase during embryonic development. Three lines of evidence support growing extrusion activity. (i) While TADs (quantified by the strength of insulation) remain weak at 5.3 hpf, they become more pronounced at later stages (ED Fig. 1c). (ii) Analysis of the contact probability P(s) curves provide complementary evidence as they can reveal the presence of loops, irrespective of barrier elements. Extruded loops create a hump on the P(s) curve, best seen in its log-derivative ^34,35^. We observe dramatically different slopes of P(s) before and after ZGA, with signatures of ∼50kb loops starting to emerge as early as 5.3hpf, yet becoming pronounced and larger (∼100-150kb) at 11 hpf (ED Fig. 1d). (iii) Published protein abundance data show presence of Rad21 cohesin component in the chromatin fraction as early as 5.3 hpf ^27^ and its growth with time. Together, this suggests that cohesin-driven loop extrusion is initiated at ZGA, and strengthens through development. Furthermore, this early loading of cohesin is consistent with our observation of fountains upon ZGA (Fig. 1b,c), if fountains are formed due to localized cohesin loading. Indeed, fountains (“jets”) in another system ^29^ were associated with cohesin loading. Functionally, early cohesin loading may be required for mediation of enhancer-promoter interactions ^16–20^.

### Fountains are transient structures emerging after ZGA

Fountains emerge at 5.3 hpf, likely representing early stages of chromosome folding. We define a fountain as a *pattern of contacts that emanates from a single genomic locus and broadens with distance from the diagonal* (Fig. 1b,c). To systematically characterize fountains, we developed an automatic *cooltools*-based ^36^ algorithm, *fontanka,* that is able to distinguish fountains from potential genomic misassemblies (ED Fig. 1e-f, see Methods). Efficient fountain calling allowed to see both individual patterns and their characteristic average shape: Fountains represent enrichments of contacts in an approximately 200 kb range from the base, comparable in size yet very different in shape from TADs. In contrast to TADs, fountains are pinched near the diagonal, have a distinct base, and lack sharp boundaries (Fig. 1c,d).

Fountains are clearly distinct structures, different from TADs. Fountains do not correspond to either TAD centers nor TAD borders (Fig. 1d, Fig. 2a, also see SI). They do not colocalize with CTCFs or sites of insulation (Fig. 2a, ED Fig. 2c) and look distinct from stripes that emanate from CTCF sites 45 degrees from the diagonal ^37^.

**Figure 2.**
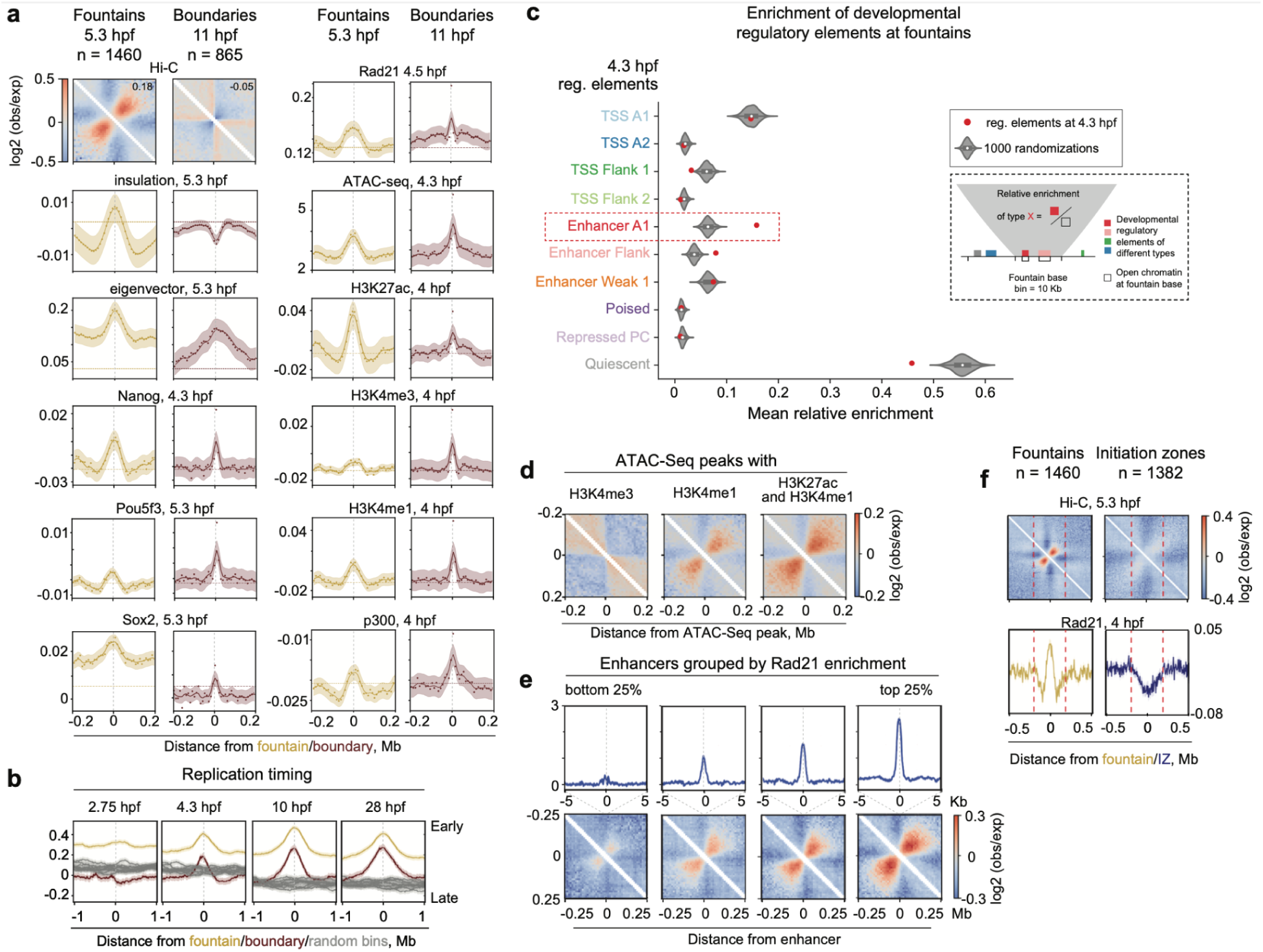
Functional characterization of fountains. **a.** Genomic features of fountain bases compared to TAD boundaries. Units: insulation score ^36,66^; Nanog^67^, Pou5f3^25^, Sox2^22^, p300^25^, and histone modifications^25^: log2 ChIP-Seq over input; ATAC-Seq^40^: reads. Note that insulation is negative at the TAD boundaries (meaning fewer average interactions) and positive at fountains (more average interactions). **b.** Fountain bases replicate earlier than TAD boundaries. Average replication time of fountain bases and TAD boundaries at four indicated developmental stages (Repli-seq data from ^38^). **c.** Fountains colocalize with enhancers at 4.3 hpf. Dashed inset: calculation of relative enrichment of regulatory element at fountains as fraction of fountain genomic bin covered by regulatory element of specific type (example for Enhancer A1). Developmental regulatory elements are predicted ATAC-seq-supported developmental regulatory elements (PADREs) from ^39^. Briefly, PADREs are ATAC-Seq at the dome stage annotated by ten ChromHMM states obtained for corresponding stage of development. We calculated the enrichment of developmental regulatory elements at genomic bins with fountains, and compared it to the control (1000 randomizations of fountain positions). Significantly enriched over control: Enhancer A1 (FDR<< 1e-4, obs. ∼0.158, exp. mean ∼0.065, exp. std ∼0.009), Enhancer Flank (FDR << 1e-4, obs. ∼0.079, exp. mean ∼0.038, exp. std ∼0.007, FDR is based on normal distribution fit, see Methods “Enrichment of developmental regulatory elements at fountains”). **d.** Fountains form at enhancers but not on promoters. Average pileups of Hi-C maps at 5.3 hpf for regions enriched in promoter (H3K4me3, H3K4me1) and enhancer (H3K4me1, H3K27ac) histone marks (also see ED Fig. 3a-b, d-e, control: ED Fig. 3g-h). **e.** Fountain strength at enhancers reflects cohesin occupancy. 5422 active enhancers (dome regulatory elements Enhancer A1, data from ^39^) were sorted to four groups by descending Rad-21 ChIP-seq signal (data from ^4^). Top panel: cohesin occupancy profiles. Bottom panel: average pileups of the enhancers. Note that fountain strength goes down with cohesin occupancy. Compare to ED Fig. 2b. **f.** Comparison of fountain bases (n=1460) with replication Initiation Zones at 4.3 hpf (IZs, data from ^38^). Only IZs closest to fountain bases were taken into account (n=1382). Red dotted lines demarcate +/- 0.2 Mb from summit). From top to bottom: (*top*) Hi-C signal, 5.3 hpf. Note that specific fountain structure forms on fountain bases but not on IZs. (*bottom*) Rad 21 occupancy, 4 hpf ^4^, on fountains versus replication Initiation Zones. Note that fountain bases are enriched in cohesin (Rad 21), while IZs are depleted of cohesin. See also ED Fig. 2d,i.

Remarkably, using *fontanka,* we detected 1460 fountains consistent across two Hi-C replicates at 5.3 hpf (Suppl. Dataset 1), many more than seen in adult mouse thymocytes (38 jets ^29^) or other systems ^28,30^ (see SI). Since fountain calling methods produce similar results (fountain score and “protractor” introduced by Guo et al. to detect “jets” ^29^, ED Fig. 1g, see also SI), such striking difference in the number of fountains between two vertebrate systems likely reflects the difference in biology between them, consistent with massive cohesin loading upon ZGA.

We next examined how fountains change through development. We observe that fountains gradually transform into other patterns at 25 hpf, such as dots and TADs, which are known to be formed by loop extrusion (Fig. 1b,c). Fountains are nevertheless visible at 11 and 25 hpf as an average pattern but hardly detectable individually, possibly masked by TADs, interrupted by CTCFs, or suppressed by other mechanisms (Fig. 1d, see also SI).

Next, we asked whether fountains are present during early development in other species. We analyzed Hi-C data for development of medaka fish ^9^ and *Xenopus tropicalis* ^6^. Using our fountain caller, we identified individual fountains at stages close to ZGA in both species, revealing a clear fountain signal similar to that observed in zebrafish (Fig. 1g,h, Suppl. Dataset 2).

Together, this analysis shows that fountains constitute patterns distinct from known hallmarks of chromosome organization: TADs, stripes, and compartments. Fountains are not unique to zebrafish, as they are also visible in other externally developing early embryos of medaka and *Xenopus*.

### Fountains are enriched at enhancers

Aiming to understand the functional role of fountains and their relationship to genome activation, we examined enrichment of genomic features at and around fountains. For comparison, we performed the same analysis for TAD boundaries that become visible at 11 hpf (Suppl. Dataset 3).

Both fountains and TAD boundaries were enriched in regions of open chromatin, active chromatin marks, cohesin, Pol II and pioneer TF binding as early as 4-5.3 hpf (Fig. 2a, ED Fig. 2a-c). Most of the fountains are located in A compartment at 5.3 hpf (65.6% of fountains versus 47.8% expected by chance), and are predisposed to avoid the future boundaries (0.2% versus 1.1% expected by chance). We also compared the replication timing between the fountain bases and boundaries, using the Repli-seq data at 2.75, 4.3, 10 and 28 hpf ^38^. We found that, consistent with their euchromatic location, fountains were replicating significantly earlier than boundaries in all time points (Fig. 2b, ED Fig. 2d).

However, fountains showed strong enrichment of enhancer-specific mark H3K4me1 and H3K27ac, and rather weak enrichment of the promoter-specific H3K4me3 mark. TAD boundaries, in contrast, did not display this preference for enhancer marks and were about equally enriched in H3K4me1 and H3K4me3 (Fig. 2a, see also SI). Together, this suggests that fountains are enriched in enhancer marks and depleted in promoter marks showing patterns distinct from that of TAD boundaries.

To further characterize the functional states of fountains, we examined the enrichment of developmental regulatory elements at 4.3 hpf (ATAC-Seq peaks annotated by ChromHMM chromatin states in development ^39^). Such functional categories include promoters of different levels of activity, active enhancers, Polycomb repressed promoters, heterochromatin, and quiescent chromatin. Surprisingly, we found that among ten developmental regulatory element types, only the enhancer states show significant enrichment at fountains, as compared to a random control (FDR<<1e-4 based on normal distribution fit, Fig. 2c). Neither transcriptionally active states, such as transcription start sites, nor inactive or repressed states are enriched at fountains as compared to a randomized control (see Methods). We found that 854 fountains are associated with sites of active transcription (ATAC-seq peaks with marks of enhancers or promoters, located within 10 Kb distance from the fountain bin, with 670 expected in the random control), with 625 fountains containing at least one enhancer (with 342 expected in the random control).

As enhancers tend to be located close to their target promoters, we asked whether fountains were proximal to active promoters. We found that fountains were indeed significantly enriched in the proximity of zygotic genes that began transcription between 3 and 5.5 hpf (p-value∼10^-8^, ED Fig. 2f, see also SI), with about ∼50% of fountains having an active gene less than 200 Kb away; and ∼70% (1042) fountains containing promoters active at dome stage within a 50 Kb window.

To further explore connection between fountains and enhancers, we performed a complementary analysis that does not rely on *a priori* identification of fountains. We stratified all ATAC-seq peaks by their enhancer (H3K4me1) and promoter (H3K4me3) marks and compared average Hi-C pile-ups centered at ATAC-seq peaks from different categories (Fig. 2d, ED Fig. 3a,b). We observed striking differences in Hi-C patterns on promoters and enhancers. Enhancers showed a pronounced fountain pattern that became even stronger when active enhancers were selected using a combination of high H3K4me1 and high H3K27ac (Fig. 2d, ED Fig. 3d,e, control: ED Fig. 3g,h). On the contrary, Hi-C pile-ups centered at promoters showed a pattern opposite to that of a fountain, i.e. an average insulation (Fig. 2d, detailed analysis in ED Fig. 3). This analysis, independent of fountain calling, complements the observation of enhancer enrichmentat at fountains and demonstrates that fountains form preferentially at enhancers rather than promoters.

Our developmental timeline allows to examine the relationship between fountains formed upon ZGA (5.3 hpf) and enhancers activated later in development (12 hpf). We found that 5.3 hpf fountains are also enriched in 12 hpf enhancers (ED Fig. 2e). Moreover, when we considered later enhancers alongside early ones, 765 fountains contained at least one such enhancer (vs 625 containing 5.3 hpf enhancers). This suggests that at least some fountains are formed at sites that will establish their enhancer status later in development.

### Fountains form at cohesin-enriched sites within early replicating regions

In zebrafish, DNA replication timing program is progressively refined in parallel with lengthening of the cell cycle in development ^38^. As fountains and non-random replication initiation zones (IZs) are established within the same developmental time window, we inquired if fountains mark the early replication forks initiating on IZs (Suppl. Dataset 4). However, the averaged Hi-C map of IZs ^38^ does not resemble the average fountain (Fig. 2f) and the fountain strength does not depend on the distance to IZs (ED Fig. 2i). Aiming to investigate the possible relationship of fountains to replication, we compared the genomic properties of fountain bases and IZs. Both IZs and fountain bases were enriched in open chromatin and enhancer marks (ED Fig. 2d), consistent with the location of fountains in active (and hence early-replicating) chromatin. However, and strikingly, fountain bases were enriched in cohesin, while IZs were rather depleted of it (Fig. 2f, ED Fig. 2d). Taken together, this suggested that cohesin accumulation on enhancers results in fountain formation independently of replication initiation.

### Cohesin accumulation is associated with fountain strength

Overall one of the most distinct features of fountains (Fig. 2) was the accumulation of cohesin. We examined this connection by two approaches, one that uses called fountains and the other that doesn’t. Among identified fountains, those with higher cohesin (Rad21 subunit) levels showed a stronger fountain pattern (ED Fig. 2b). Independent of fountain calling, we grouped strong enhancers by cohesin levels and found that higher cohesin signal correlated with stronger fountains pattern (Fig. 2e). Together, this indicates that fountains preferentially form at cohesin-rich enhancers. Perturbation experiments below allow to test a causal connection between enhancer activation and establishment of fountains.

### Pioneer factors establish open chromatin platforms for fountain formation

Early establishment of enhancers in zebrafish is controlled by the binding of the key zygotic genome activators, transcription factors Pou5f3, Sox19b, and Nanog (PSN). These factors act as pioneers by creating open chromatin regions and recruiting histone modifiers, such as p300, to their binding sites ^21,22,24–26,40^. If fountain formation requires chromatin accessibility and enhancer activity, we expect the removal of Pou5f3, Sox19b or Nanog to disrupt fountain formation on the regions where these factors bind and open chromatin. To test this mechanism, we performed Hi-C on the 5.3 hpf embryos, null-mutant for Pou5f3 ^41^, Sox19b ^40^, Nanog ^42^ and also on double or triple mutant embryos bearing combinations of two or three of these mutations (MZ*triple*) (Fig. 3a).

**Figure 3.**
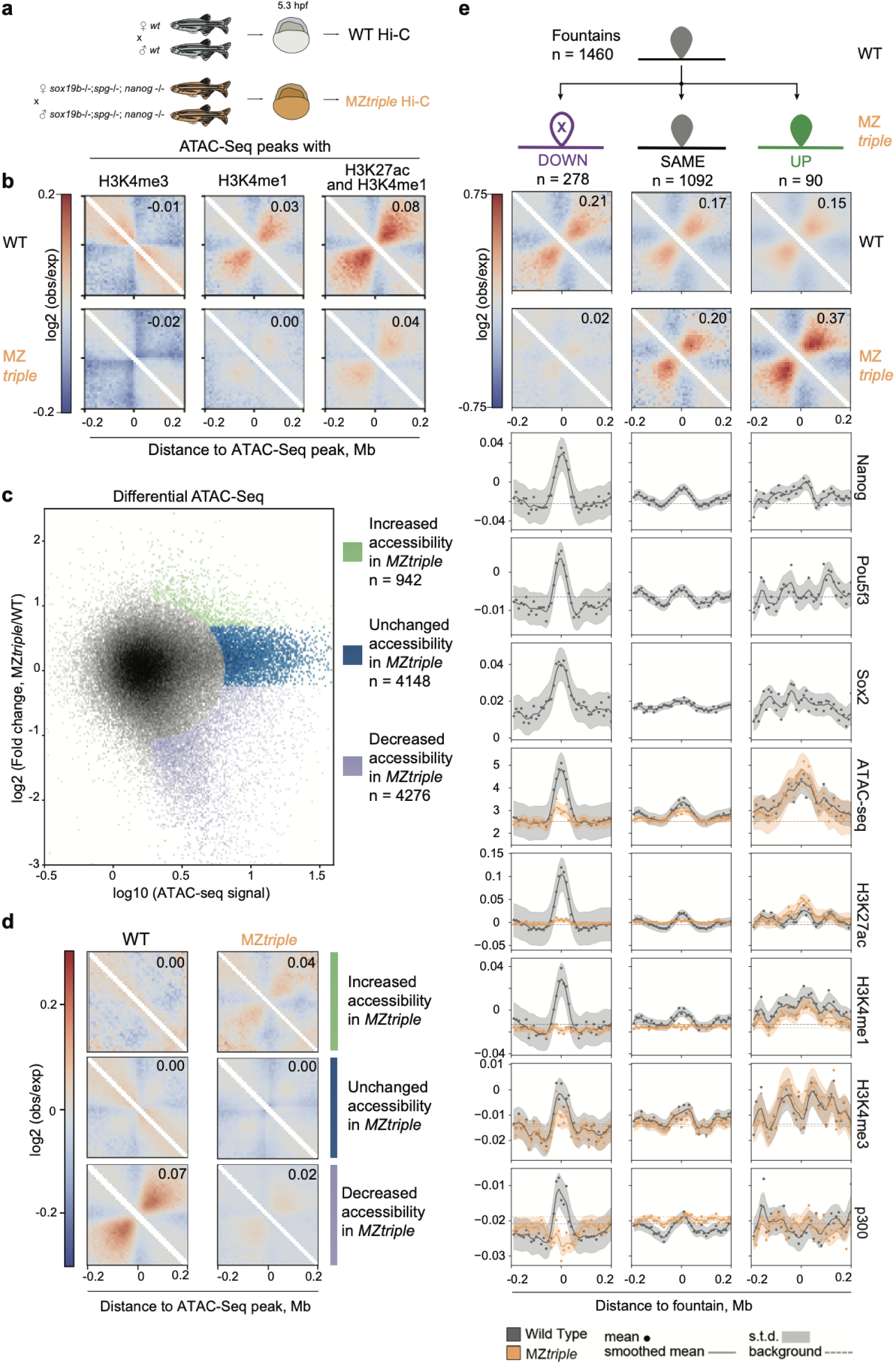
Pioneer factors establish open chromatin platforms for fountain formation (three approaches). WT – wild type; MZ*triple* – Maternal-Zygotic (MZ) homozygous Pou5f3/Nanog/Sox19b mutant. Numbers in the top corners of average pileups represent average fountain strength for the pile of snippets. **a.** Schematics of obtaining triple Pou5f3, Sox19b or Nanog mutant of zebrafish to test the formation of fountains. See Methods and Riesle et al. 2023 ^43^ for details. **b.** Abolishment of fountain signature in MZ*triple* at the ATAC-Seq peaks enriched with different type of active histone marks, similar to Fig. 2d. See ED Fig. 3a for detailed explanation of the stratification design. (**c-d**). Chromatin openness in MZ*triple* mutant is coupled with fountain pattern formation on Hi-C. **c.** Distribution of ATAC-seq responses in MZ*triple* compared to wild type and definition of the groups of genomic bins based on their ATAC-seq responses. Data from ^40,43^. **d.** An average pileup at ATAC-Seq peaks from different groups from (c) in wild type Hi-C and MZ*triple* Hi-C at 5.3 hpf. **e.** Weakened fountains in the triple Pou5f3/Nanog/Sox19b (MZ*triple*) mutant bear the strongest early enhancer features. From top to bottom: (*top*) Three groups of fountains by the change in fountain strength in MZ*triple*. Out of 1460 fountains, 278 weakened (DOWN), 90 strengthen (UP), and 1092 remained unchanged (SAME, see Methods “Differential fountains in MZ*triple*” for definition). (*middle*) Pileups of wild type and MZ*triple* Hi-C at DOWN-, SAME-, and UP-fountains. (*bottom*) Enrichment profiles of various features of chromatin at three groups of fountains. ATAC-Seq: mean read count; other profiles: log2(ChIP-Seq/input). ChIP-seq for Nanog at 4.3 hpf from ^67^ and for Pou5f3 and Sox19b at 5.3 hpf from ^25^, ATAC-seq data from ^25^. H3K27ac and p300 ChIP-seq data for 4 hpf from ^25^.

First, we examined changes in Hi-C patterns at four classes of ATAC-seq sites: enhancers, promoters, both, and neither (Fig. 3b). Fountain pattern, seen most distinctly for the enhancer group, has vanished in MZ*triple*, strongly supporting the role of enhancers in establishing fountains.

Next, we examined sites of open chromatin (ATAC-seq peaks) that changed in MZ*triple* (Fig. 3c). We found that sites that lost their accessibility in MZ*triple* showed a fountain pattern in the wild-type that vanished in MZ*triple* (Fig. 3d, also see SI). Similar effects were seen for accessible sites with either H3K27ac or H3K4me1 marks (ED Fig. 3j-k).

Finally, we performed a differential analysis of the identified fountains (Fig. 3e) and categorized them into three groups: fountains that weakened in MZ*triple* (Down, n=278), remained unchanged (Unchanged, n=1092), and became stronger in MZ*triple* (Up, n=90). The weakened fountains showed the strongest enhancer signature and were the most enriched for pioneer transcription factors binding (PSN, Fig. 3e). However, many fountains formed at this stage remain unaffected, suggesting that they are either controlled by PSN-independent enhancers or by enhancers that become active at later stages of development.

However, many fountains formed at this stage remain unaffected, suggesting that they are controlled by PSN-independent enhancers. Below we show some of these PSN-independent enhancers become fully active later in normal development, accompanied by the strengthening of fountains at them.

Together, this triple-knockout has revealed that PSN-dependent enhancers active at ZGA form fountains. Moreover, abolition of their enhancer activity led to the loss of the fountain signature, indicating that enhancer activity is required for fountain formation at these sites.

Finally, we demonstrate that even individual pioneer TFs contribute to fountains formation as seen from changes of the fountain strength for individual Pou5f3 ^41^ and Nanog ^42^ mutants. Recently we found that Pou5f3 and Nanog promote chromatin accessibility in different genomic locations ^43^, and, thus, we used chromatin accessibility as a proxy to the Pou5f3-, Nanog- and Pou5f3/Nanog-dependent enhancers. In total, 441 out of 1460 fountain bases lose their accessibility in any of the mutants (30% of all fountains with FDR <5%, Supplemental dataset 5). More thorough classification of these fountains revealed that TF-specific loss of chromatin accessibility led to specific loss of fountains (ED Fig. 4). This analysis further strengthens the causal link between establishment of accessible chromatin by pioneer TFs and formation of fountains.

### Fountains are formed at functional sites that are established later in development

Since most of the fountains remained unchanged and some even got stronger in MZ*triple*, we asked next what distinguishes these fountains from the PSN-dependent ones that weaken in MZtriple mutant. Indeed fountains that strengthen in MZ*triple* also got stronger later in normal development, after gastrulation (11 hpf, Fig. 4a). They also became earlier replicating after gastrulation (Fig. 4b), and acquired H3K27ac after gastrulation (12 hpf, Fig. 4c). These observations echoed our recent finding ^43^ that late enhancers become aberrantly open and activated at ZGA in MZtriple.

**Figure 4.**
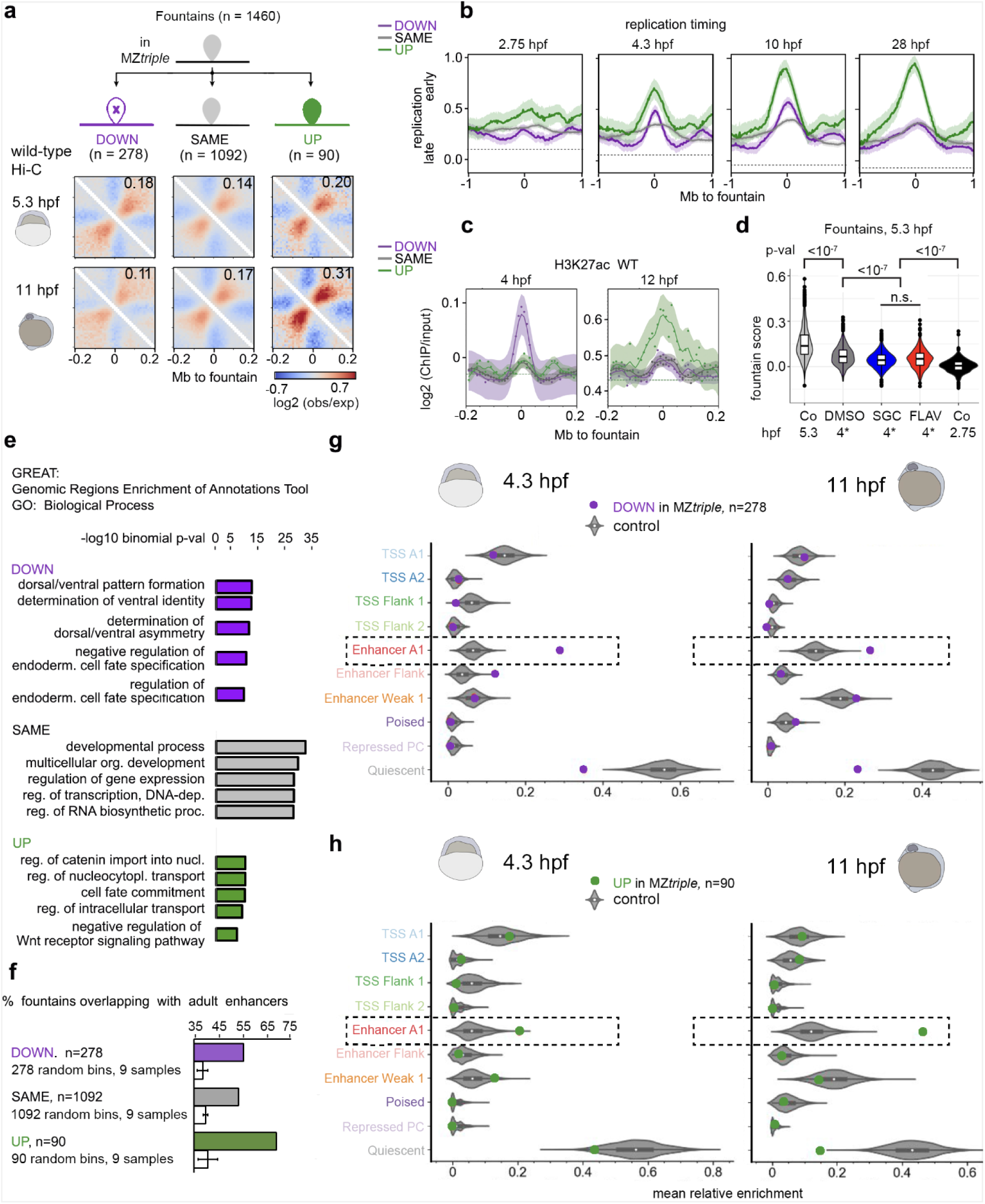
Changes in enhancer repertoire underlie fountain changes in MZ*triple* mutant. **a.** Fountains upregulated in MZ*triple* mutant at 5.3 hpf (“UP”) increase later in normal development. Top: three groups of fountains by regulation in MZ*triple,* the color code is used throughout this figure. Bottom. Hi-C average fountains at 5.3 hpf (WT2, two replicates, see Suppl. Dataset 3) and at 11 hpf. **B**. Replication timing on three groups of fountains (“UP”, “SAME”, DOWN”) during developmental progression. **c.** H3K27ac on three groups of fountains (“UP”, “SAME”, DOWN”) at 4 hpf and 12 hpf. **d.** Fountain strength is reduced in the embryos treated by p300 and RNApolII inhibitors (datasets marked by 4* hpf are from ^4^). Violine plots show fountain scores for 1460 fountains in the indicated conditions. DMSO – control, SGC – p300 inhibitor; FLAV – RNApolII inhibitor, p-values in Tukey-Kramer test. **e.** Top 5 GREAT enrichment categories in GO: Biological Process for “DOWN”, SAME” and “UP” fountains. **f.** Percentage of fountains overlaping with enhancers from 11 adult zebrafish tissues (data from ^45^). Fountain (or control bin) was scored as overlapping if it overlapped with at least one enhancer for at least 1 bp. Bin size is 10 kb, error bars represent standard deviation. **g.** Fountains weakened in MZ*triple* mutant (DOWN) colocalize with early (4.3 hpf) enhancers. Developmental regulatory elements data from ^39^. Analysis similar to Figure 2c. **h.** Fountains strengthened in MZ*triple* mutant (UP) colocalize with late (12 hpf) enhancers. Developmental regulatory elements data from ^39^. Analysis similar to Figure 2c.

Interestingly, the reduction of H3K27ac upon p300 inhibition or blocking of the RNApolII by flavoperidol at 4 hpf ^4^ resulted in some weakening of fountains but not in their collapse (Fig. 4d). This observation suggests that key to fountain formation are processes upstream of transcription initiation and H3K27 acetylation such as TF binding and the establishment of accessible chromatin on enhancers.

Majority of fountains (*∼*75%) that were unchanged in MZ*triple,* and thus are PSN-independent, may nevertheless be enriched in weaker functional sites and/or enhancers that get activated later in development. PSN-independent fountains were enriched in active enhancer marks, but less than the fountains of two other categories (Fig. 4c). Using functional annotation of regulatory sites ^44^, we found that PSN-independent fountains are enriched in general developmental regulatory categories (Fig. 4e). Similarly to PSN-dependent groups, PSN-independent fountains were enriched in overlapping tissue-specific adult enhancers ^45^ (Fig. 4f).

Consistently, fountains that go down in MZ*triple* correspond to early active enhancers, i.e., had ATAC-seq peaks with active enhancer signature at 4.3 hpf (Fig. 4g). Fountains that go up in MZ*triple* correspond to enhancers active after gastrulation, i.e., were enriched with active enhancer signature at 12 hpf (Fig. 4h). Altogether, our analysis established a causal relationship between the changes of chromatin accessibility on enhancers and fountain formation.

### Facilitated loading of loop extruding SMCs аs a mechanism of fountains formation

We argue that, while distinct from TADs and stripes, fountains reflect early chromosome folding by cohesin-mediated loop extrusion. First, we found that extruded loops of ∼50 kb emerge as early as 5.3 hpf (ED Fig. 1d). Second, we observe cohesin accumulation at fountains at this stage (Fig. 2a, ED Fig. 2a-c). Finally, fountains with high cohesin levels show a more pronounced pattern of contacts (ED Fig. 2b).

Known extrusion-mediated patterns are formed by stopping loop extrusion at specific positions, resulting in the accumulation of cohesin at these positions ^37,46^. Fountains are distinct from TADs and stripes yet also accumulate cohesin at their bases (Fig. 2a, ED Fig. 2a-c). Since we see no enrichment of CTCF at fountains (ED Fig. 2c), cohesin accumulation there can be due to another mechanism. In fact, the strength of fountain shape demonstrates no dependence on the distance to extrusion barriers, such as CTCFs and actively transcribing genes (ED Fig. 2g-i). Thus, we hypothesize that cohesin accumulation at fountain bases, and fountains themselves, emerge due to facilitated cohesin loading.

To test the facilitated loading mechanism, we developed polymer simulations in which cohesin loads preferentially at specific sites (Fig. 5a). We defined the site of facilitated cohesin loading as a 1-kb region (one simulation monomer), consistent with recent characterizations of regulatory elements ^47^ and the average length of ATAC-seq peaks at fountain bases (606 bp) ^39^, which corresponds to the narrow (∼10 kb) bases of the fountains we observe. Once loaded (facilitated or uniform), cohesins perform two-sided loop extrusion, stopping when they encounter one another ^12^.

**Figure 5.**
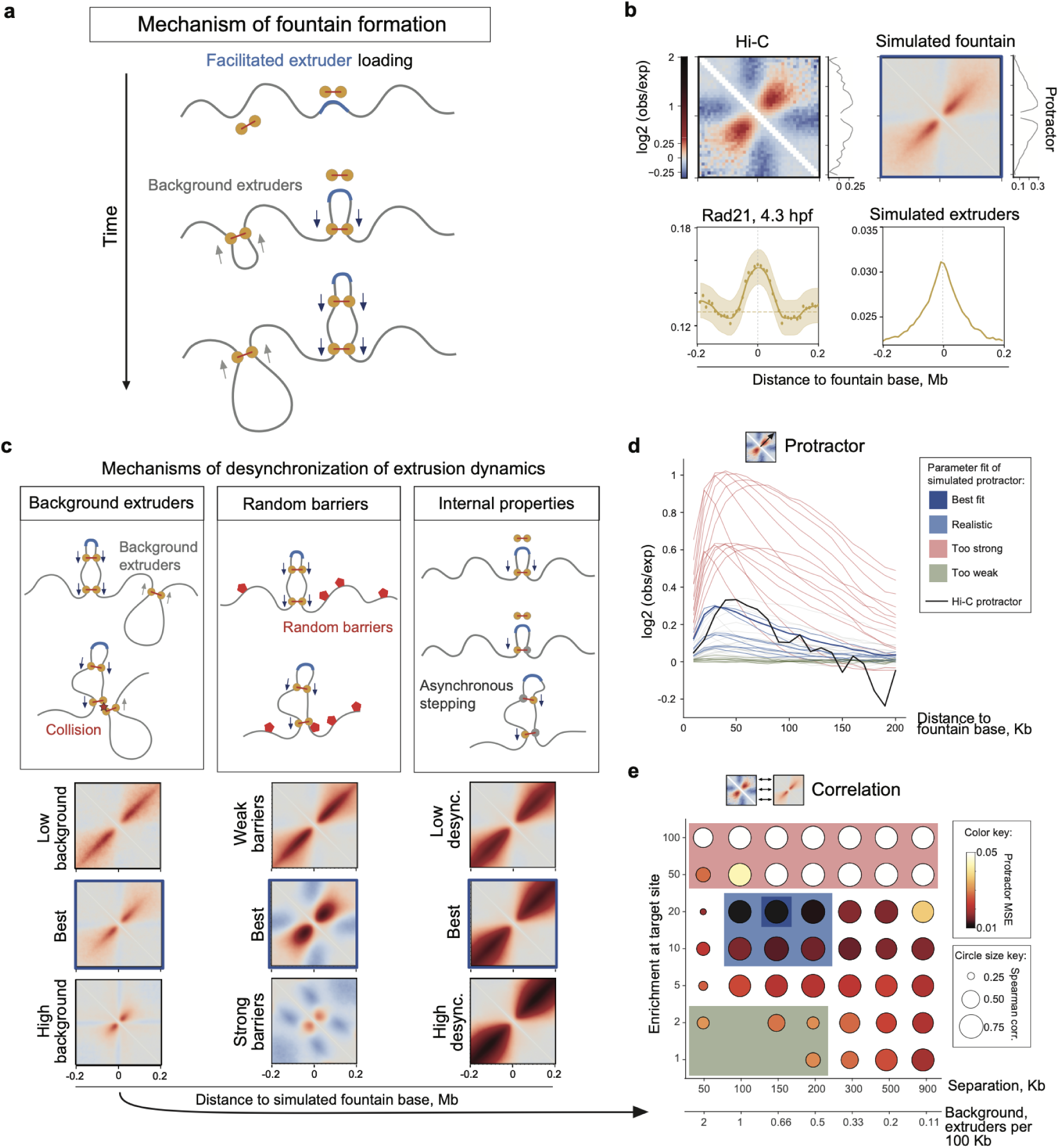
Facilitated extruder loading аs a mechanism of fountains formation. **a.** Proposed mechanism of fountain formation. The extruder molecule (cohesin complex) loads preferentially at the target region (highlighted in blue) and starts extruding DNA in two directions. Multiple extruders can load at the target sites, resulting in the formation of a hairpin-like structure. Extruder molecules can also load at other genomic loci (background extruders with lower rates of loading). **b.** Comparison of Hi-C average fountain to the best fit of the model with facilitated loading and background cohesins. (*top left*) Average Hi-C pileup based on experimental data, log2 observed-over-expected signal. (*bottom left*) Distribution of cohesin subunit Rad21 around 5.3 hpf fountains, log2 ChIP-Seq over input signal at 4.3 hpf ^4^. (*top right*) Average simulated Hi-C pileup based on simulations with the best parameters according to (**d-e**, also see Methods “Goodness of fit of the simulations to real data”). (*bottom right*) Cohesin coverage profile in simulations, calculated as the number of extruders per simulation monomer accumulated throughout simulation normalized by the simulation size. **c.** Three mechanisms of desynchronization of extrusion dynamics: (*left*) cohesin movement is desynchronized due to collisions with background cohesins, (*center*) cohesin sides are randomly stopped by random barriers, (*right*) internal properties of cohesin lead to asynchronous stepping of two sides. All three scenarios lead to emergence of fountains with increasing spread with distance from the loading site. (**d-e**). Parameter sweep of the model of facilitated cohesin loading in the presence of background cohesin loading (**c**, *left*). **d.** Protractor measurement of simulated fountains for average fountain from Hi-C (black) and various simulations parameters. Color scheme as in (**e**): red – fountains are too strong, green – fountains are too weak, blue – fountains are in the similar range to the Hi-C data. Protractors are displayed for the same parameters as in (**e**). **e.** Similarity of the simulated fountains to the real Hi-C data. Each circle represents simulation round and comparison of *in silico* average fountain to the Hi-C fountain (ground truth). The color represents mean square error (MSE) of protractor characterising the intensity of interactions at the fountain. The size of circles represents the Spearman correlation of the fountain shape (in ±200 Kb-snippets). Boxes of different colors represent various types of simulations outcomes, color scheme as in (**d**).

### Fountain shape helps elucidate loop extrusion mechanism

Next, we identified conditions and mechanisms that lead to the formation of fountains in simulations. We observed that the loading of extruders only at the loading site and extrusion from there yields not a fountain, but rather a narrow hairpin pattern similar to that observed in bacteria (ED Fig. 5a-c). Gradual broadening of experimental fountains (Fig. 5b) as they emanate from the fountain bases suggests that the extrusion path of cohesin is not perfectly symmetric, i.e., reeling of DNA on one side of cohesin is not perfectly synchronized with another side. Asymmetric extrusion by cohesin, when averaged over cells, could produce symmetric fountains with increasing spread with distance from the loading site as we measured in Hi-C (Fig. 5c, ED Fig. 5a-c).

We suggest three mechanisms that can lead to such asymmetry in extrusion dynamics (Fig. 5c). The first mechanism is the intrinsic asymmetry of extrusion by cohesin, i.e. one-sided extrusion with direction switching, as was long anticipated ^37^ and recently observed in vitro ^48^. The second mechanism is external to the motor and constitutes pausing of enhancer-loaded cohesins on randomly positioned barriers: while stochastically pausing at a barrier, cohesin can continue reeling chromatin on the other side. Such randomly positioned extrusion barriers can be molecules of MCMs ^49^ or transcribed genes ^46^. The third mechanism is the pausing of enhancer-loaded cohesins at other cohesins. Such mechanism stipulates that cohesin loads uniformly along the genome, yet with a preferential loading at enhancers. This mechanism can also explain the weakening of fountains later in development due to the rising levels of cohesin on DNA ^27^.

For every model we swept its parameters and selected models that best fit shapes of fountains from Hi-C for 5.3 hpf (Fig. 5c, ED Fig. 5,6). All models have a range of parameters where they could closely approximate experimental fountains. For the model of background and facilitated loading, our parameter sweeps (Fig. 5d-e, ED Fig. 5a-c) revealed that ∼10-fold facilitation at narrow (1 Kb) enhancer sites is sufficient to closely match the intensity profile of fountains. As for the background cohesins, the density of 1 cohesin per 100-200 Kb is required, consistent with cohesin density in other vertebrate systems ^12^. Simulations of this mechanism reproduce the correlation of the fountain shape (Fig. 5e), intensity of signal at fountains (measured by protractor tool, Fig. 5d) and cross-sections of fountains at the genomic distances up to 200 Kb (ED Fig. 5c). Finally, simulations also reproduce a broad (∼50-100 Kb) peak of cohesin accumulation seen in ChIP-Seq (Fig. 5b).

**Figure 6.**
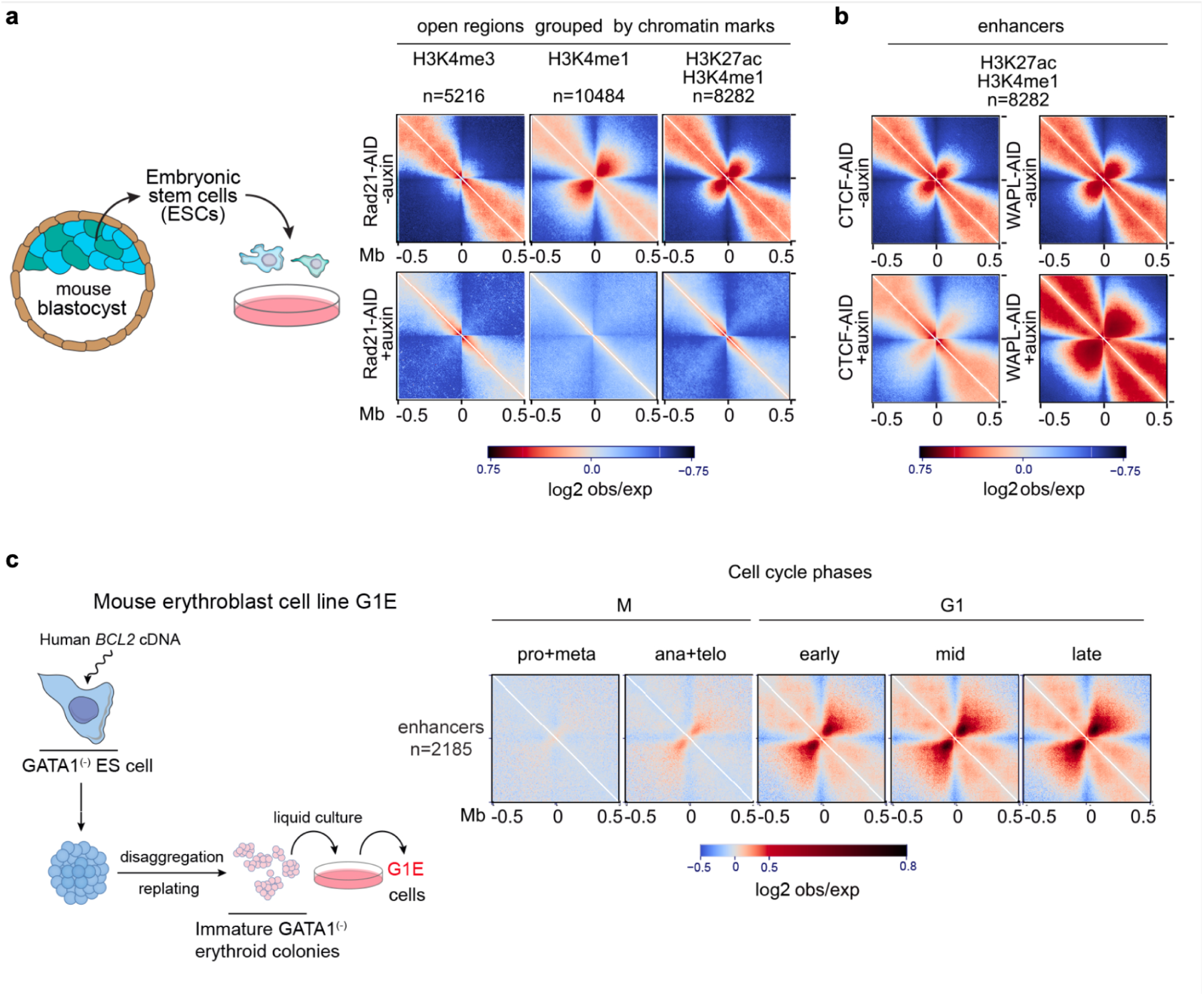
Cohesin-dependent fountain-like structures emerge on the enhancers in mouse cell lines. **(a-b).** Average Micro-C pileups at the indicated genomic elements in mouse ES cells in two conditions. Top: control condition (no auxin), bottom: auxin-inducible acute knockout of the indicated gene (Micro-C from ^56^). Open chromatin regions were assessed by DNase sites from ^68^, and histone modifications – by ChIP-Seq from the same study. Overlaps with CTCF peaks were excluded from the analysis in all cases (CTCF ChIP-Seq data from ^56^, see Methods “Hi-C snipping and average pileup”), bin size 5 Kb. **a.** Fountains form on enhancers and depend on cohesin in mouse ES cells. Left: Embryonic origin of mouse ES cells. Right: Pile up on non-CTCF open chromatin regions (DNAse I narrow peaks) grouped by enrichment on indicated chromatin marks for mouse embryonic stem cells E14 (ENCODE consortium data ^69^) in control and upon Rad21 acute degradation. Note that fountain-like structures form on enhancers but not promoters (compare to Fig. 2d), bin size 5 Kb. **b.** Pile up on enhancers (defined as H3K4me1+ H3K27ac+ open chromatin regions, same as in Fig. 6a, right) in control and upon CTCF and WAPL acute degradation. **c.** Fountain-like structures on non-CTCF potential enhancers form during G-phase and start to emerge as early as in ana-telophase. Left: Origin of G1E erythroblast cell line; modified from ^70^. Right: Pileup on active enhancers in G1E erythroblast cell line synchronized in the indicated phases of the cell cycle (Hi-C data and ChIP-Seq from ^57^). Overlaps with CTCF peaks were excluded from the analysis in all cases. Examples of individual fountains formed at enhancers in G1E cells can be found in ED Fig. 7 and SI Fig. 12.

We also considered and ruled out an alternative mechanism where fountains are formed by affinities of fountain-proximal regions, i.e., compartment-like mechanisms. Such a mechanism would result in enrichment of contacts between fountains that we don’t see in the data (ED Fig. 5d).

Importantly, all extrusion-based mechanisms do not rely on CTCF for the formation or broadening of fountains, consistent with the absence of CTCF near fountains in zebrafish (ED Fig. 2c). Consistently, a recent study by Isiaka et al. ^30^ observed prominent fountains in *C. elegans* that lack CTCF (lost in nematode evolution). This contrasts the formation of “stripes” at CTCF-bound sites across vertebrate genomes ^12,50^. Interestingly, in simulations, each fountain contains, on average, ∼1 cohesin in the steady state. This indicates that they are different from bacterial hairpins, where multiple (∼15-30) extruders of bacterial SMC were simultaneously moving along a hairpin from the loading to the unloading sites ^51,52^. Together, our modeling suggests that fountains can be formed by one of these mechanisms or their combination: facilitated loading, interactions between cohesin and with other extrusion obstacles, or internally desynchronized extrusion of cohesin motors.

### Cohesin-dependent fountain-like structures emerge at enhancers in mouse cells

To elucidate further the role of cohesin in generating fountains, we asked whether fountains are present in early development of mammalian species, and if their formation depends on cohesin. Uniquely, mouse systems allow to study the effect of cohesin loss due its experimental depletion as well as natural cohesin loss during metaphase and its reloading upon entrance into G1.

In mouse embryonic stem cells (mESC), a model for inner cell mass of mouse preimplantation blastocyst, pioneer factors Pou5f1 and Sox2 mediate cohesin binding on enhancers ^53–55^, in striking parallel to our zebrafish system. We examined Mirco-C from mESC, as well as in lines where acute cohesin, CTCF and Wapl depletions were performed ^56^. Although we could not automatically call individual fountains as they were overshadowed by TADs, dots and stripes, we piled Micro-C maps at CTCF-distal enhancers and promoters (see SI). In the control condition, we observed a distinct fountain pattern on enhancers but not on promoters, as we observed in fish (Fig. 6a, top, compare to Fig. 2d). Strikingly, enhancer-associated fountain pattern vanished upon cohesin degradation (Fig. 6a, bottom). We also see changes in fountain shapes upon CTCF degradation (Fig. 6b, bottom), further supporting cohesin as an extruder underlying fountain formation. The enhancer-associated fountain pattern is more concentrated in untreated cells, and becomes more extended upon CTCF degradation. This is consistent with CTCFs stopping enhancer-loaded cohesin and preventing their spreading further away. Moreover, Wapl depletion that leads to increased cohesin density and processivity shows much stronger fountains. Taken together, this analysis establishes a strong causal link between cohesin-driven extrusion and the formation of fountains at enhancers during the early stages of embryonic development.

If fountains are formed due to cohesin-mediated extrusion we expect them to vanish during mitosis and gradually reappear as cohesin reloads on chromosomes in early G1. To this end, we examined fountains in Hi-C data from cell cycle-synchronized populations of cultured G1E erythroblast cells ^57^. Consistent with cohesin’s role, fountains on piled enhancers were absent during prometaphase and metaphase, and began to reappear upon exit from mitosis, gradually strengthening from early to mid to late G1 phase (Fig. 6c, ED Fig. 7). These results support the cohesin-mediated mechanism for fountain formation and further indicate that this process occurs independently of replication.

## Discussion

By analyzing chromosome organization in early zebrafish development, we found that loop extrusion activity initiates upon ZGA. Yet structures formed by extrusion are different from those seen in adult cells such as TADs and dots. A new class of chromosome structures – fountains, with more than a thousand emerging across the genome around the time of genome activation. We demonstrated that fountains are distinct from known features of local chromosome folding, like TADs, dots, and stripes—all of which are linked to the pausing of extruding cohesin by CTCFs ^12^, and to a lesser extent by transcribed genes ^46^. Fountains, in contrast, are not associated with CTCFs or positions of TAD borders, and do not directly depend on transcription. Importantly, we demonstrated that fountains are not just a peculiarity of a Hi-C map, but rather, they show a striking association with developmental enhancers and causal relationship with them. In synergy with our efforts, studies of Isiaka et al.^30^ and Kim et al. ^31^ found enhancer-associated fountains (jets) in *C. elegans*, the organism which lacks CTCF, supporting our finding that fountains are formed on enhancers by a CTCF-independent mechanism.

Association of fountains with enhancers is two-way specific, i.e., among ten types of functional annotations, active enhancers were the only ones enriched at locations of fountains. Conversely, an average Hi-C map at enhancers shows a pronounced fountain pattern. Importantly, the latter observation did not rely on definitions or calling of the fountains.

Knockouts of key pioneer transcription factors establish a causal link between early active enhancers, chromatin accessibility, and fountains. Knockouts of individual or combinations of transcription factors result in the loss of chromatin accessibility and enhancer activity at specific locations, leading to the corresponding loss of fountains at those sites. Interestingly, only 19% of fountains are affected by the loss of all three pioneer transcription factors, suggesting that other genomic elements may also contribute to their formation.

Our analysis suggests that fountains also form at latent enhancers, which become active only later in development and in specific tissues. In zebrafish, regulatory elements show little overlap between tissues and developmental stages ^39,45^. These late and tissue-specific enhancers do not carry specific enhancer marks at 5.3 hpf and cannot be readily discerned. However, we found that fountains unaffected by the loss of pioneer transcription factors are enriched in enhancers that become active after gastrulation and in adult tissues, and show a broad association with developmental genes. Altogether, this suggests that fountains formed upon ZGA form on active as well as on latent enhancers.

Which factors enable fountain formation at active and latent enhancers? The most parsimonious explanation is chromatin opening by pioneer factors. Many of such factors are already present in the embryo at sufficiently early stages, e.g. zebrafish orthologs of mouse pioneer transcription factors Eomesodermin and Brachyury ^58,59^. These factors can establish accessible chromatin at latent enhancers, thus triggering fountain formation, way before these enhancers are fully activated. While enhancer activity is necessary for fountain formation, fountains are not essential for formation of enhancers – a conclusion of a recent study in adult *C.elegans* ^30^ where loss of fountains upon cohesin COH-1 cleavage did not result in enhancer loss. Indeed, some histone marks and accessibility are mitotically inherited ^60,61^, while others quickly established in G1, allowing rapid fountain formation in early G1 as we observe here (Fig. 6c).

Several lines of evidence show association of fountains with cohesin. First, we found accumulation of cohesin at fountains, with the higher levels of cohesin associated with stronger fountains. Similarly, enhancers with higher levels of cohesin show a stronger fountain pattern. Second, fountains at enhancers in mouse cells vanish upon acute cohesin (Rad21) depletion, while getting much stronger upon depletion of Wapl, when cohesin processivity and density are increased. Furthermore, the loss of fountains in mitotic cells and their gradual reemergence in G1 upon cohesin reloading are consistent with the cohesin-mediated mechanism. Consistently, a double knockout of CTCF and Wapl in mice reported formation of “plumes” at open chromatin regions ^28^. A recent study in *C.elegans* further shows fountain loss upon acute cohesin depletion ^30,54^. Another recent study of partial depletion of cohesin *in vivo*, in postmitotic mouse dendritic cells, reported weakening of fountains that is accompanied by drastic loss of immune function of these cells ^20^. Finally, our simulations and those of Guo et al.^29^, demonstrate that facilitated loading of cohesin followed by bidirectional extrusion leads to formations of the fountain pattern. Importantly, our simulations also reveal that (i) about ∼10 fold facilitation is sufficient, and that (ii) such facilitated loading needs to be accompanied by mechanisms that desynchronize reeling of DNA into an extruded loop from the two sides of cohesin. Such desynchronization can be an inherent property of cohesins, as was seen *in vitro* ^48^, or can result from collisions with randomly positioned barriers and/or with other extruding cohesins elsewhere.

While it is natural to expect cohesin loading at accessible regions, such as enhancers, surprisingly, other accessible regions, such as promoters, do not show any association with fountains. In fact, active promoters show an “anti-fountain” insulation pattern, consistent with the role of transcription as an extrusion barrier ^46^. Given the overall similarity between enhancers and promoters, selective loading of cohesin on enhancers is surprising. Two mechanisms could lead to the formation of fountains selectively on enhancers. The first mechanism would rely on selective recruitment of cohesin components, e.g. by pioneer transcription factors, to enhancers as have been reported for mouse ES cells ^53–55^, human cancer cell lines ^62^, mouse liver ^63^ and cultured *Drosophila* cells ^64,65^. Second mechanism requires a factor that would interfere with cohesin bidirectional movement at promoters, such as moving polymerase ^46^, rendering fountains indistinguishable specifically at promoters.

Fountains are readily observable in zebrafish embryos, but they can be hard to discern in adult cells, with the only reported instances being in quiescent thymocytes ^29^, and early in the M-to-G1 transition (ED Fig. 7). Although facilitated loading at enhancers persists throughout development and the cell cycle, the distinct shapes of individual fountains become less apparent. Indeed, collisions between facilitated-loaded cohesin and other cohesins or polymerases—as well as stalling and sequestration at CTCFs—transform fountains and overshadow them with TADs and other structural features. Consistent with this mechanism, we find that at early developmental stages, cohesin accumulates at fountains rather than at CTCF sites, whereas at later stages, cohesin redistributes from fountain bases to CTCF sites and TAD borders (ED Fig. 2c). Similarly, fountains appear at enhancers in the mouse cell cycle as early as anaphase/telophase (Fig. 6c, ED Fig. 7), but become increasingly difficult to distinguish from dots and stripes at later stages of mitotic exit (ED Fig. 7). The continued detection of fountain signatures at enhancers in later developmental stages and during later interphase indicates that the facilitated loading process remains active. Altogether, these observations suggest that fountains are most prominent in early development and early G1, eventually evolving into well-recognized extrusion-mediated structures.

While fountains facilitate contacts between regions flanking an enhancer, they do not facilitate contacts of the enhancer itself. This prompts the question: how can fountains help enhancers to perform their functions? One possibility is that facilitated cohesin loading at enhancers facilitates contacts between other regulatory regions on one side of an enhancer with genes on the other, thus helping other nearby enhancers. Another possible role of enhancer-loaded cohesin can be to stall at enhancer, thus performing a one-sided extrusion, and thus facilitate formation of its interactions with distal genomic elements. Such loading of cohesins at enhancers may be part of the mechanism by which cohesin’s loop extrusion facilitates enhancer-promoter interactions ^16–20^. If stalling at enhancers and one-sided extrusion events are sufficiently infrequent, they might not leave a noticeable signature in the fountain pattern, yet facilitate contacts with target genes. Ultimately, facilitated loading of cohesin at enhancers constitutes a promising avenue for future investigation into the principles governing enhancer–promoter communication.

Broadly, our analysis shows that loop extrusion activity initiates upon ZGA. Yet structures formed by extrusion change through development due to different activity of cohesin-regulating factors. Cohesin-loading activity appears to be the primary factor mediating chromosome folding upon ZGA, while cohesin-stopping factors shape chromosomes at later stages. Surprisingly, key regulatory sites driving transcription, early enhancers, also appear to be the elements driving early folding events by recruiting cohesins. Extruding from enhancer, cohesins form the new elements of chromosome architecture – fountains. This new role of enhancers in actively folding chromosomes can shed new light on the mechanisms of long-range regulatory activity of enhancers in development and beyond.

## Data availability

Raw and processed Hi-C data for zebrafish embryogenesis is available at GEO at https://www.ncbi.nlm.nih.gov/geo/query/acc.cgi?acc=GSE195609. The list of zebrafish fountains is available in Suppl. Dataset 1, Xenopus and medaka fountains in Suppl. Dataset 2, zebrafish TAD boundaries in Suppl. Dataset 3, zebrafish initiation zones at 4.3 hpf in Suppl. Dataset 4, and fountains group assignment based on chromatin accessibility change in mutants is available in Suppl. Dataset 5. The detailed list of the sequencing datasets is available in Suppl. Dataset 6.

## Code availability

*Cooltools*^36^-based tool *fontanka* for fountain calling with an example is available at https://github.com/agalitsyna/fontanka with the code of fountain calling under *examples* directory. Simulations of facilitated loop extrusion are available at https://github.com/agalitsyna/polychrom_workbench under *targeted_extrusion* directory.

## Author contributions

SU: Hi-C libraries; AxG: Hi-C and genomic analysis, polymer simulations; MB: replication timing data analysis, NB, KP, MiG: bioinformatics; SU, MV, MeG, DO: zebrafish mutants; DO: study design; SR, DO: supervision of the wet part, funding acquisition; MiG, LM: supervision of Hi-C genomic analysis and simulations. AxG wrote the first draft, and SU, SR, LM, and DO developed and edited the manuscript.

## Acknowledgments

We are grateful to Edward Banigan for proofreading the text and to all members of the Mirny lab for many productive discussions. We thank Damir Baranasic and Chris Sansam for sharing unpublished data. AxG thanks Max Imakaev, Nezar Abdennur, and Anton Goloborodko for early ideas for the polymer simulations; Henrik Pinholt, Simon Grosse-Holz, and Emily Navarrete for ideas on the model and figures; Antoine Coulon, Mark Pownall, and Christopher Bohrer for the thorough discussions of the results and critical assessment of the hypotheses; students Alexey Shkolikov (FBB MSU) and Eli Rybnikova (PRIMES MIT) for their preliminary analysis of fountains. DO is grateful to Prof. Lev Yampolsky for the consultation about statistical analysis. This work was supported by DFG-ON86/5-1 and DFG-EXC2189 for DO, by GM114190 to LM, and by Russian Science Foundation (grant #21-64-00001) to SR.

---

**Extended Data Figure 1.**
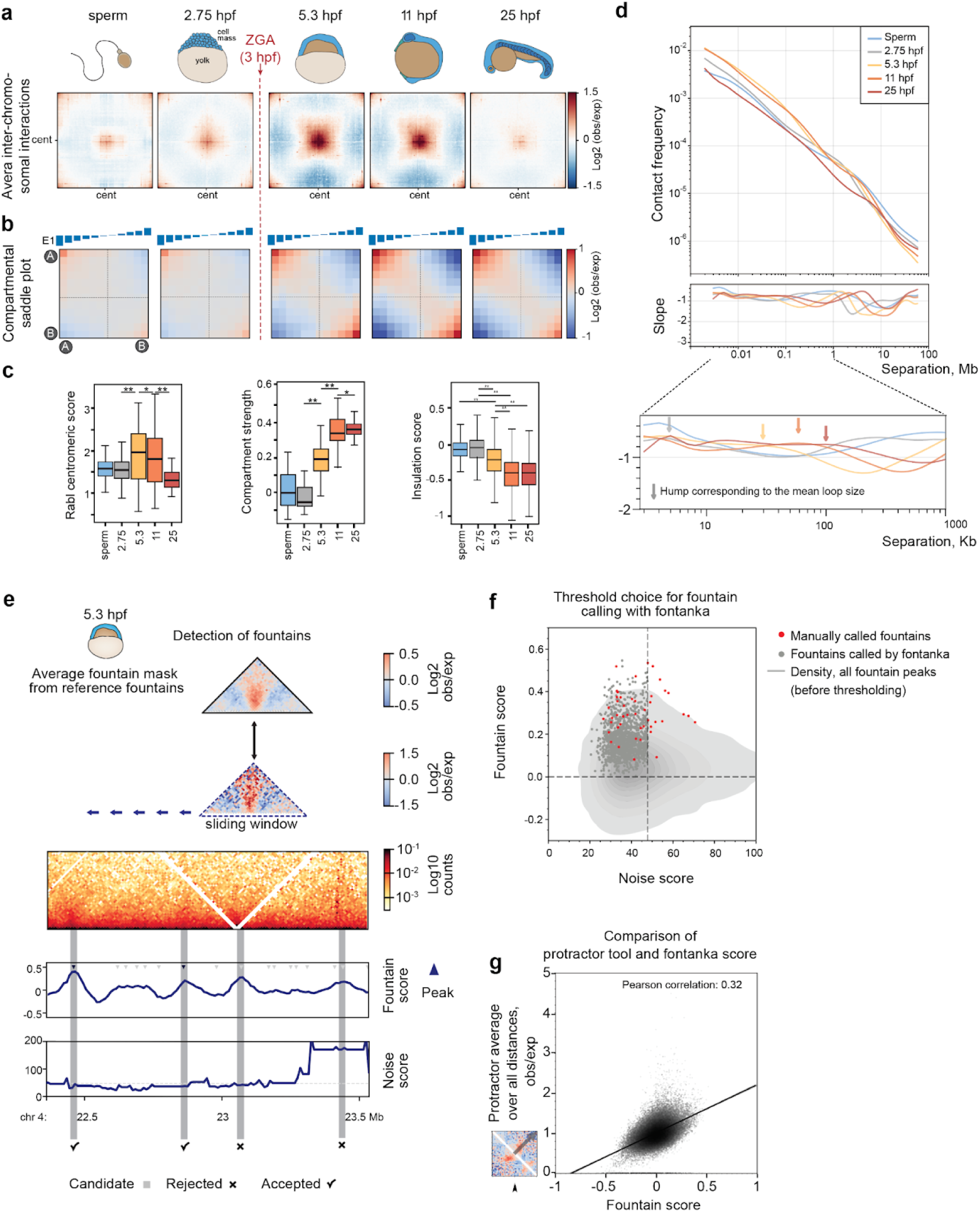
Establishment of 3D chromatin in development, finding of fountain structures. (Supporting material for main Fig. 1). **a.** Centromere-centromere interactions measured by average Rabl plots. **b.** Compartmentalization measured by saddle plots. **c.** Boxplot of Rabl centromeric scores (*left*), compartment strength (*center*), and insulation score (*right*) at different stages of embryo development. Insulation score: sperm and 2.75 hpf boxplots are based on boundaries found at the same stage; 5.3, 11, and 25 hpf insulation are based on boundaries found at 25 hpf. Note that insulation is usually negative at the TAD boundaries (meaning fewer average interactions). **d.** Scaling plots for different stages of development of zebrafish. From top to bottom: (*top*) Dependence of the contact probability *P*_c_(*s*) on the genomic distance *s* for sperm and the four indicated developmental stages. (*middle and bottom*) First derivative indicating the slope of *P*_c_(*s*). The region between 3 kb and 1 Mb is additionally magnified (*bottom*). Arrows indicate the approximate positioning of the scaling humps, which are detectable as broad elevations of interactions rather than clear local maxima. Note that the short-range peak for 2.75 hpf is almost absent, while it starts to be more discernible at 5.3 hpf and shifts toward larger genomic separations. The peak reaches 100 Kb typical for other Hi-C datasets of vertebrates ^34^ at 25 hpf. **e.** Detection of fountains with the fountain-calling algorithm *fontanka*. log2 observed/expected ratios of manually picked fountains at 5.3 hpf were averaged to obtain a fountain mask (*top*). The fountain mask was compared to a 0.4 Mb sliding window to obtain the fountain score for each 10 kb genomic bin (*second from top*). The peaks in the fountain score (*second from bottom,* gray triangles) were considered fountains if their noise score was smaller than the threshold (*bottom*, see Methods for full fountains selection criteria). The 1460 fountains found on the 5.3 hpf contact map using this procedure are referred to as “fountains” and are further analyzed in this work. **f.** Setting up the thresholds of fountain peak score and noise (Scharr) score for selecting fountains, see Methods “Fountain calling with *fontanka*”. **g.** Application of protractor tool ^29^ for validation of fountains. The grey arrow in the inset (fountain) represents the perpendicular to the main diagonal (ideal hairpin). The scatterplot shows the correlation of the protractor tool score ^29^ with the fountain score defined in this paper. See Methods “Comparison of fontanka and protractor tool” and SI for average protractor for fountains.

**Extended Data Figure 2.**
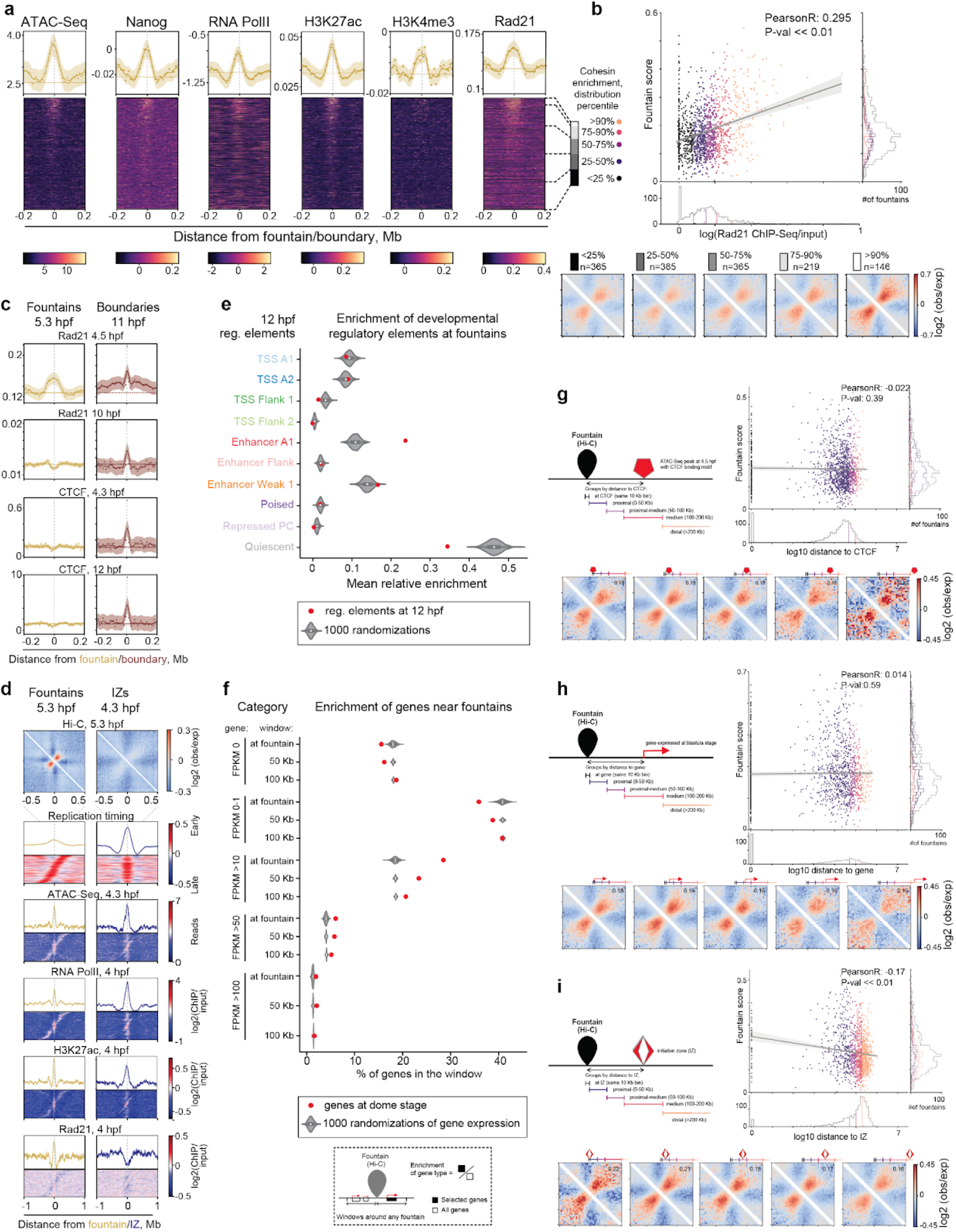
Properties of the fountains. (Supporting material for main Fig. 2). **a.** Profiles and heatmaps of the factors associated with fountains are shown for ChIP-seq data for Rad21 ^4^, RNA polymerase II ^25^, Nanog ^67^ and H3K27ac ^25^, and ATAC-seq data ^40^ at the indicated stages. The heatmaps were sorted by descending fountain score, as indicated on the right. Note that all factors positively correlate with the fountain score. Bottom: scales for the heatmaps. Units of ATAC-Seq^40^: reads; Rad21^4^, Nanog^67^, RNA PolII^25^, and histone modifications^25^: log2 ChIP-Seq over input. **b.** Fountain score correlates with cohesin occupancy. From top to bottom: (*top*) Scatterplot of 4.5 hpf Rad21 ChIP-Seq (OX) against fountain score (OY) colored by the percentiles of Rad21 signal distribution ^27^. Each dot represents a single fountain in the genome. The gray line represents linear regression on all categories. (*bottom*) Average pileup of Hi-C at 5.3 hpf for different groups with Rad21 enrichment at fountains from (*top*). **c.** Cohesin (Rad21) and CTCF occupancy on fountains before (4.5 hpf) and after gastrulation. ChIP-seq data for cohesin at 4.5 and 10 hpf from ^27^. As a proxy of CTCF occupancy, we used ATAC-seq signal on CTCF motifs at 4.3 and 12 hpf ^71^ (see Methods “CTCF binding inference from ATAC-seq”). Note that CTCF is not enriched at fountains at neither time point. **d.** Fountains are distinct from replication Initiation Zones (IZs, supporting material for main Fig. 2f). Different epigenetic marks at fountain bases (n=1460) and replication Initiation Zones (IZs 4.3 hpf, data from ^38^). Only IZs closest to fountain bases were taken into account (n=1382). Fountains were sorted by distance to the nearest IZ and vice versa. Note that both features are enriched in enhancers but only fountains are enriched in cohesin. **e.** Fountains colocalize with enhancers at 12 hpf, regulatory elements from ^39^. Similar to main Fig. 2c. **f.** Fountains preferentially localize near actively transcribed zygotic genes. We took all the genes in the neighborhood of fountains of the specified window sizes, and calculated the percentage of genes of each category (red dots). “At fountain” means window of bin size located at fountain. Control: randomly assigned categories of genes. Genes and their expressions are from EBI expression atlas ^72^. Potential maternal transcripts were excluded from the analysis (maternal transcripts from ^43^). (**g-i**). Fountain strength is independent of CTCF binding, proximity to the nearest expressed gene or replication initiation zone. Each plot, from top left to bottom right: (*top left*) Definition of the categories of fountains relative to each feature. (*top right*) Distribution of fountain strength (fountain score, OY) is related to the distance to the nearest feature (OX). Each dot is a fountain. The gray line represents linear regression on all categories. (*bottom row*) Average pileups of fountains by groups defined by distance to the nearest feature, as indicated above each pileup. **g.** Fountain strength is independent of the distance to CTCF binding site as assessed by the open chromatin region with CTCF motif (see Methods “CTCF binding inference from ATAC-seq”). **h.** Fountain strength is independent of the distance to Transcription Start Sites of expressed genes (data from EBI expression atlas ^72^). **i.** Fountain strength is independent of the distance to replication Initiation Zones (IZs 4.3 hpf, data from ^38^).

**Extended Data Figure 3.**
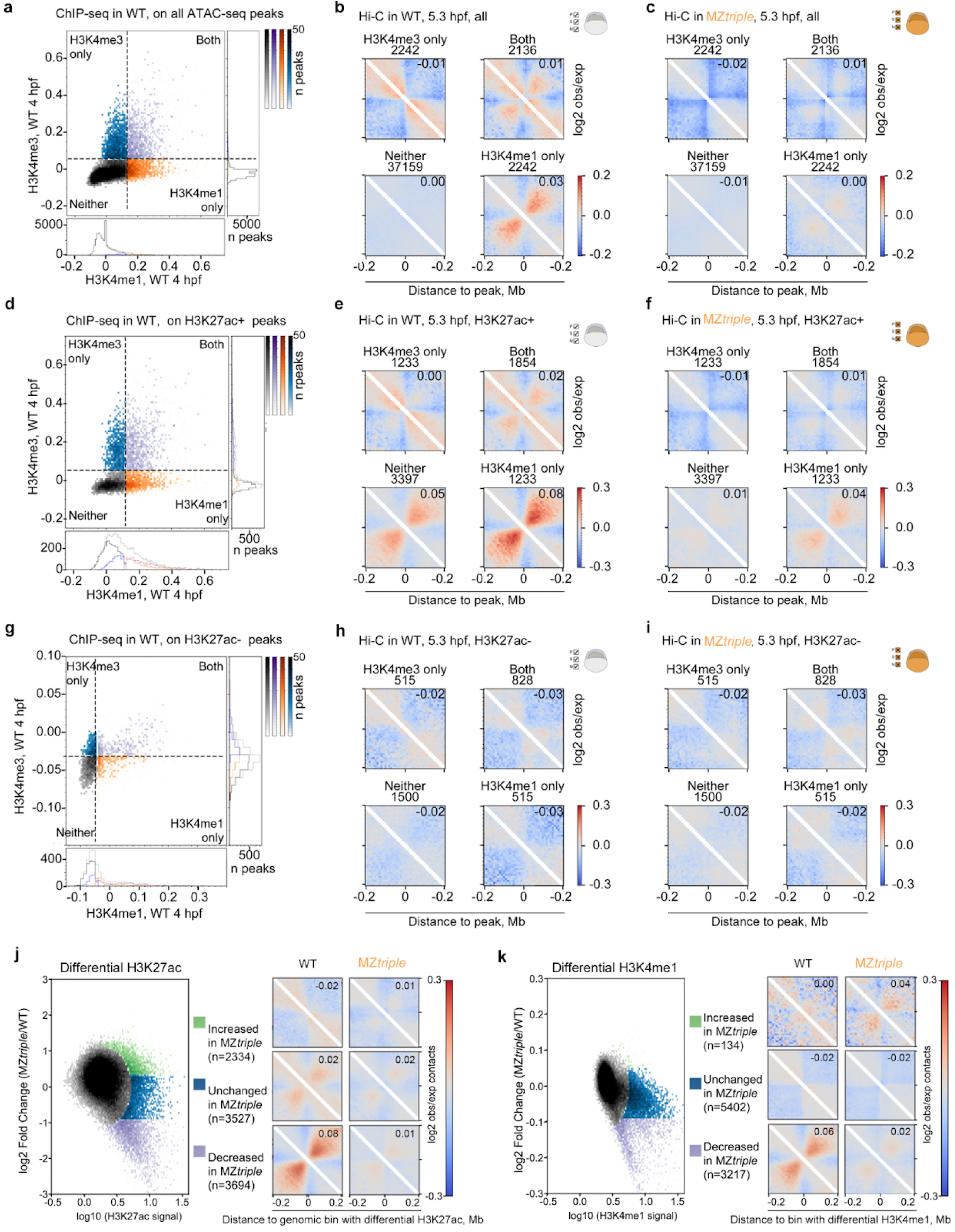
Fountains form at accessible chromatin regions enriched in H3K4me1 and H3K27ac and depend on the pioneer transcription factors Pou5f3, Sox19b, and Nanog. The average fountain score is shown at the top right corner of each Hi-C pileup. (**a-i**). Accessible chromatin regions in the wild-type embryos at 4.3 hpf (ATAC-seq peaks from ^40^) sorted by enrichment in H3K4me1, H3K4me3, and H3K27ac histone modifications at 4 hpf ^25^. Related to Main Figures 2d and 3b. Rows: (**a-c**) All ATAC-seq peaks, n = 43779. (**d-f**) Accessible regions also enriched in H3K27ac (top 10% by H3K27ac signal), n = 7717. (**g-i**) Accessible regions not enriched in H3K27ac (bottom 10% by H3K27ac signal), n = 3358, control for (d-f). Columns: (**a,d,g**). Scatterplots for the histone modifications and definition of four groups ( “Neither”, “Both”, “H3K4me1 only”, “H3K4me3 only”). Colors represent the division of the scatterplot by 90% percentile for each axis. (**b,e,h**). Average on-diagonal pileup plots for genomic regions falling into specific groups defined in (**a,d,g**), respectively, in 5.3 hpf wild-type embryos. (**b**) On all the accessible regions, fountains are detectable in the “H3K4me1 only” and “Both” groups. (**e**) On the accessible regions with high H3K27ac, the strongest fountain signal is seen on the “H3K4me1 only” group, enriched in of H3K27ac and H3K4me1 but not H3K4me3, i.e., on the active enhancers. Fountain strength increases in the order “Both” < “Neither” < “H3K4me1 only” groups. (**h**) In the absence of the H3K27ac mark, fountains do not form at the accessible regions. (**c,f,i**) Average on-diagonal pileup plots for genomic regions falling into specific groups defined in (**a,d,g**), respectively, in MZ*triple* 5.3 hpf embryos. Note that the fountain formation is absent or severely reduced in all groups, compared to the wild-type (**b,e,h**). (**j-k**). Changes in H3K27ac (**j**) and H3K4me1 (**i**) in MZ*triple* mutant are coupled with fountain pattern formation on Hi-C. Related to Main Figure 2c,d. (**j, left**) Distribution of H3K27ac responses in MZ*triple* compared to wild type and definition of the groups of genomic bins based on their H3K27ac responses. (**j, right**) An average pileup at H3K27ac bins from different groups from (j, left) in wild type Hi-C and MZ*triple* Hi-C at 5.3 hpf. (**k, left**) Distribution of H3K4me1 responses in MZ*triple* compared to wild type and definition of the groups of genomic bins based on their H34me1 responses. (**k, right**) An average pileup at H3K4me1 bins from different groups from k, left) in wild type Hi-C and MZ*triple* Hi-C at 5.3 hpf.

**Extended Data Figure 4.**
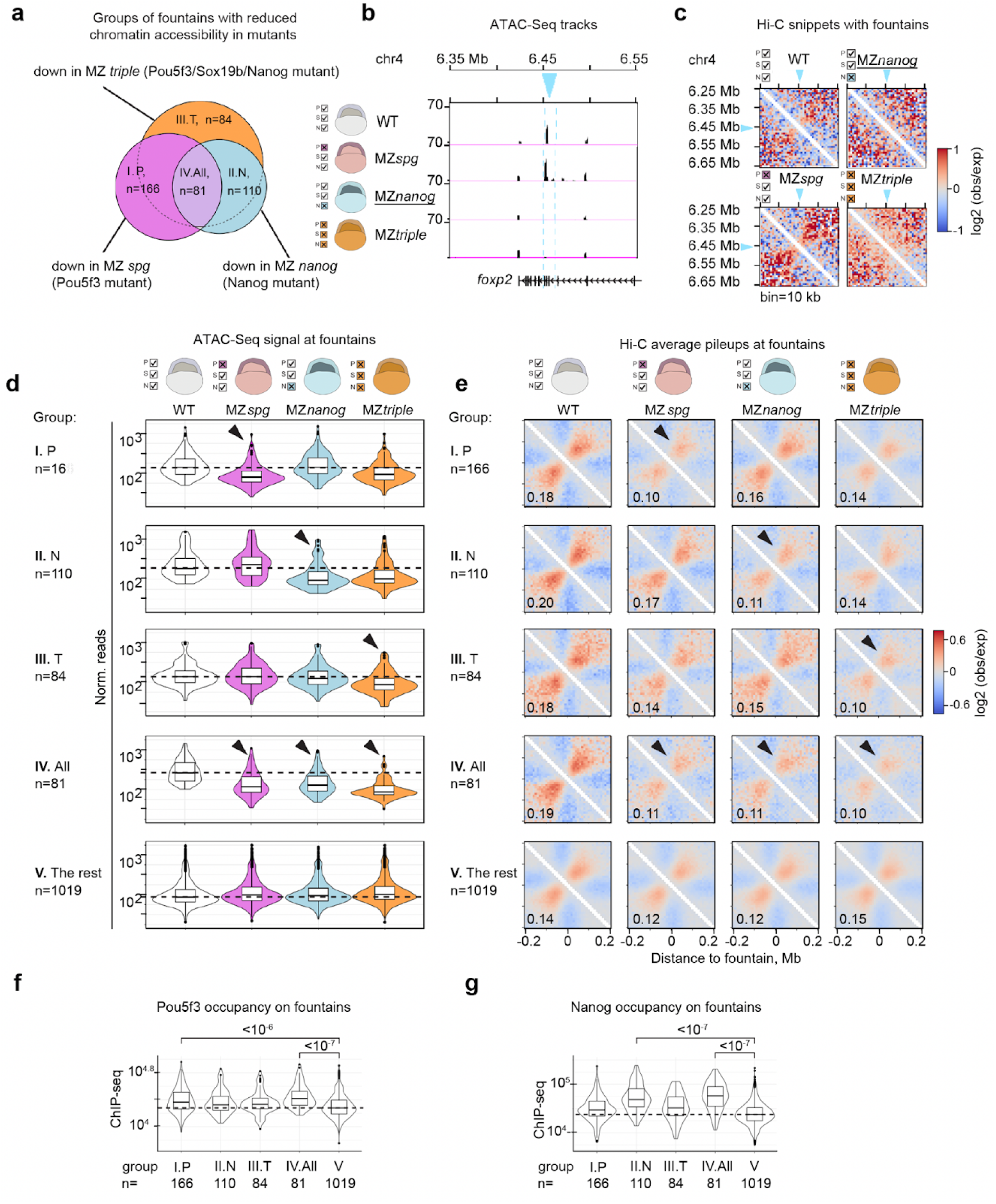
Pioneer activities of Pou5f3 and Nanog separately contribute to the formation of the fountain structures. (Supporting material for the main Fig. 5). **a.** The Venn diagram shows the groups of fountains selected by the requirement for Pou5f3, Nanog, or their combination for chromatin accessibility at the fountain bases (groups I-IV). The groups of fountains were assigned by the differential analysis of normalized ATAC-seq reads per 10 kb fountain base. ATAC-seq data for the indicated mutant genotypes and the wild-type at 4.3 hpf from ^43^. (**b, c**). Example of a fountain where both chromatin accessibility (b) and fountain structure (c) are regulated by Nanog but not by Pou5f3. **b.** The 10 kb fountain base is shown by blue arrows and dotted lines. ATAC-seq signals (rkpm) in the wild-type, Nanog mutant MZ*nanog*, Pou5f3 mutant MZ*spg* and triple mutant MZ*triple*, in 200 kb window, 4.3 hpf ATAC-seq data from ^43^. Note that in this example only Nanog is required for chromatin accessibility. **c.** Hi-C signal in 400 kb window centered at the fountain in the indicated genotypes. The fountain base is represented by the blue arrow. Note that only Nanog is required for fountain formation: the fountain is present in the wild-type and MZ*spg* but disappears in MZ*nanog* and MZ*triple*, accompanying the changes in chromatin accessibility visible in (*b*). **d.** Violin plots represent ATAC-seq signal at five groups of fountains (I-V). Groups I-IV are defined by the requirement for Pou5f3, Nanog, or their combination for chromatin accessibility, as shown in (a). Group V (the rest) contains all fountains where Pou5f3 and Nanog were not required for chromatin accessibility. Dashed lines show the median wild-type ATAC-seq signal. Arrowheads show the genotypes used to assign the groups (where the ATAC-seq signal was significantly reduced in all fountains). 4.3 hpf ATAC-seq data from ^43^. Group assignment and the corresponding ATAC-Seq signal are listed in Suppl. Dataset 5. **e.** Average on-diagonal pileup plots with log2 observed/expected ratios of the groups defined in (d). Note that the chromatin accessibility and fountain structure strength are regulated in parallel by the same factors (black arrowheads). The average fountain score is shown at the bottom left corner of each panel. (**f, g**). Pou5f3 and Nanog bind to the fountains they regulate. ChIP-seq signals for Pou5f3 (f) and Nanog (g) at 10kb fountain bases for the groups I-V. Note that each TF is enriched the most at the fountain group that it regulates: Pou5f3 at “I. P” and “III. T” groups, Nanog at “II. N” and “III. T” groups. Dashed lines show the median TF occupancy in the non-regulated group V (the lowest in both cases). P-values in the Tukey-Kramer test are shown on the graphs; p-value in 1-way ANOVA < 2e-16 in both cases.

**Extended Data Figure 5.**
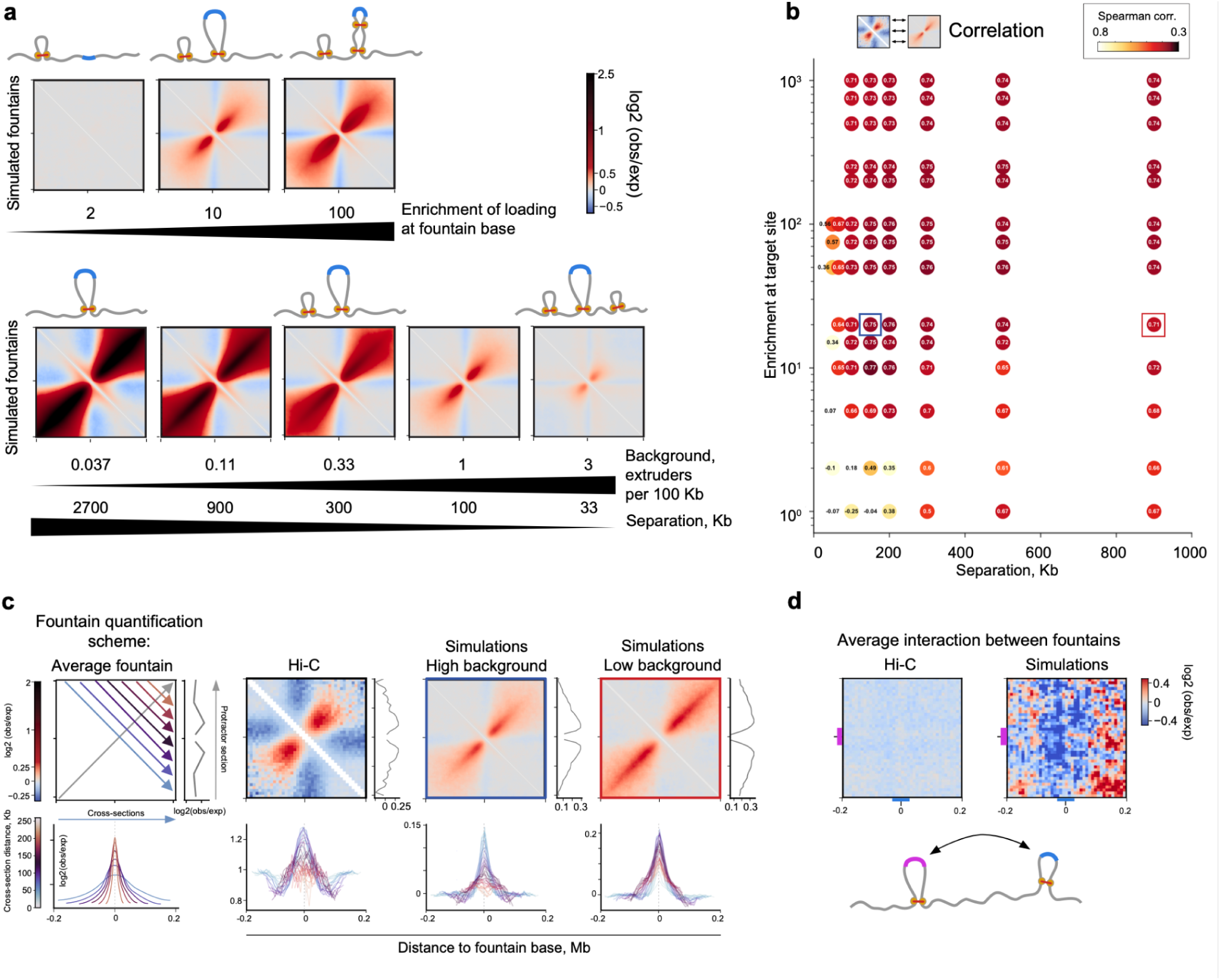
Simulations of facilitated cohesin loading. (Supporting material for the main Fig. 5). **(a-b).** Parameter sweep for the model of facilitated extruder loading with background extruders. **a.** Average fountains for simulations with different parameters of facilitated extrusion. **b.** Heatmap of Spearman correlation coefficient between Hi-C and simulated average fountains. Detailed heatmap for Figure 5e. Boxes represent the set of parameters selected for figure (**c**), blue – best parameters resulting in dispersive fountain, red – worse parameters resulting in non-dispersive fountain. **c.** Fountain quantification for real Hi-C data and two simulations from (**b**). (*left*) Scheme of quantification: calculation of protractor (*right subplot*) and cross-sections (*bottom subplot*) of average fountain. Experimental Hi-C data (*second from left*) resembles disperse fountains formed in the presence of the background loading (*third from left*) but is less similar to the non-disperse model of fountains (*right*). Difference between models can be explained as the growth of dispersion of Hi-C signal enrichment with genomic distance, noticeable at cross-sections (*bottom subplots*). Note how both Hi-C and simulated dispersive fountains cross-sections are similar in terms of broadening with distance. The parameters for simulations as in (**b**). **d.** Fountain-fountain interactions in Hi-C data (*left*) and simualtions (*right*). Absence of enriched interactions between fountains rules out compartment-like mechanism of fountain formation, where fountains could be formed by affinities of fountain-proximal regions. In the potential compartment-like mechanism, which we do not explore in the current work, two genomic regions (magenta and blue) have affinity for each other and form fountains around them. However, these “sticky” regions of different fountain bases would also generate enrichment of interactions between fountains. We examined Hi-C contacts between fountains in real Hi-C data (*left*) and found no such enrichment, ruling out the affinity-mediated mechanism. The same absence of fountain-fountain interactions can be detected in the simulations of facilitated extrusion (*right*).

**Extended Data Figure 6.**
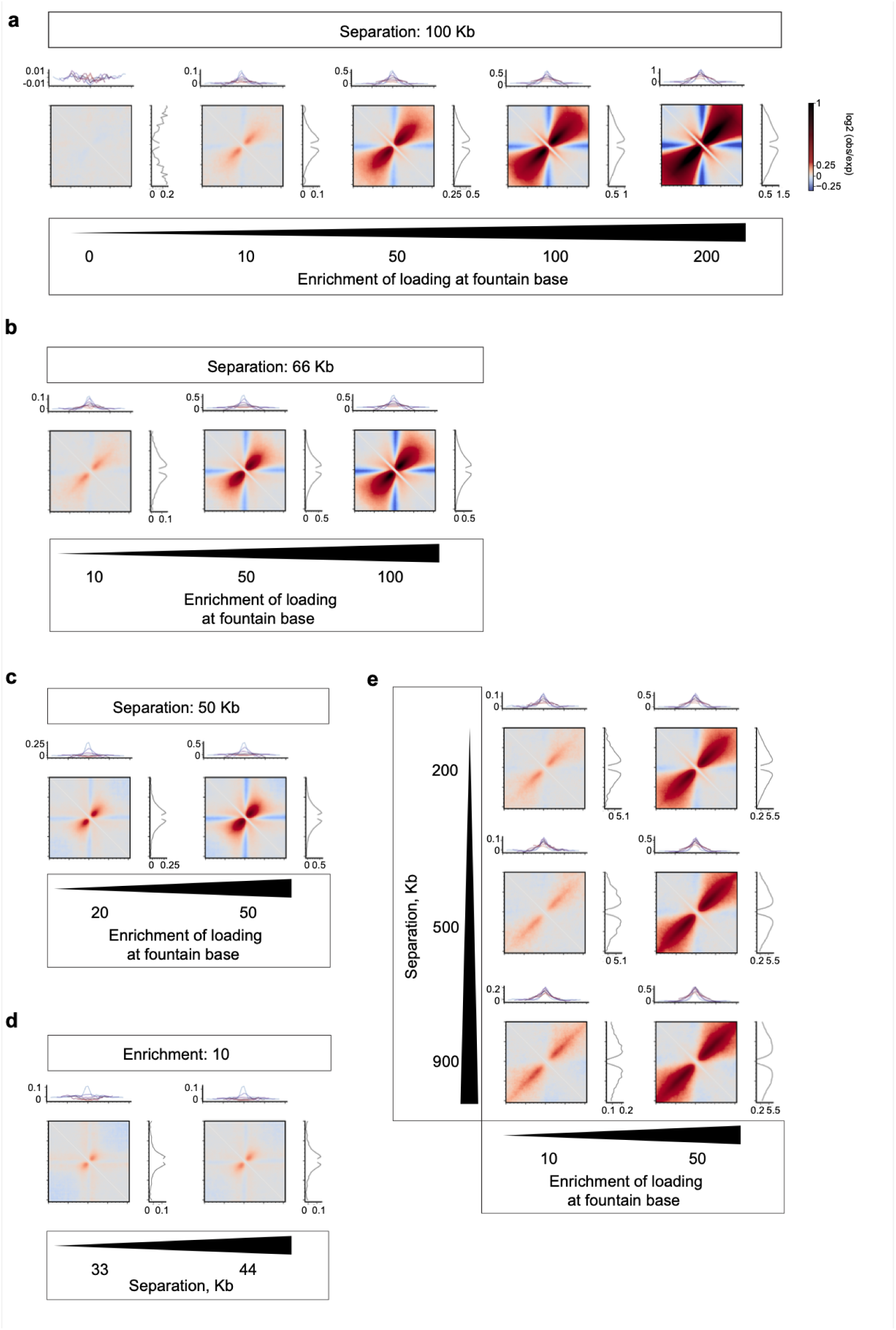
Fountain shapes in different modes of polymer simulations of fountains for the model of facilitated extruder loading with background extruders. (Supporting material for the main Fig. 5). Each plot elements: (*top*) Cross-sections of the simulated fountain as defined in ED Fig. 5c. (*center*) Simulated average fountain. (*right*) Protractor of the simulated fountain as defined in ED Fig. 5c. **a.** Simulated fountains for various enrichments at fountain base with fixed separation of 100 Kb, extended ED Fig. 5a. **b.** Simulated fountains for various enrichments at fountain base with fixed separation of 66 Kb. Note that the fountain shape changes slightly different than in (**a**), which can be attributed to more cohesins in the background than in (**a**). **c.** Simulated fountains for various enrichments at fountain base with fixed separation of 50 Kb. Note that the fountain shape changes slightly different than in (**a,b**), which can be attributed to more cohesins in the background than in (**a,b**), fountain is more suppressed. **d.** Simulated fountains for two very small separations of background extruders with fixed enrichment at the fountain base of 10, extended ED Fig. 5a. Note how the fountain is generally suppressed. Two stripes emerge from the base under smaller separations (33 Kb) due to large number of stalled extruders of the background. **c.** Simulated fountains for varying both enrichment and separation.

**Extended Data Figure 7.**
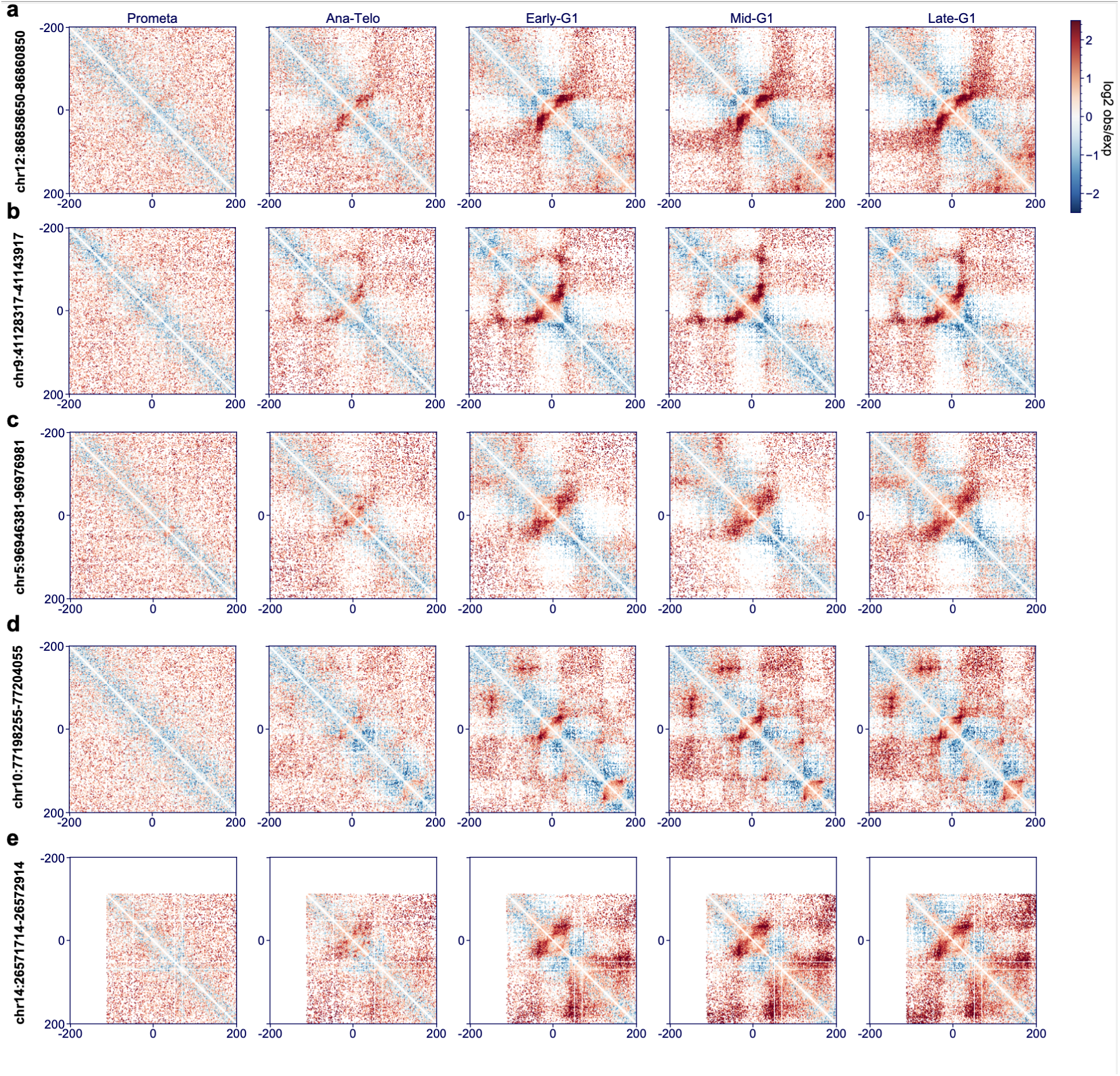
Examples of individual fountains in G1E erythroblast cell line emerging in the cell cycle. (a-e) top five fountains called based on H3K27ac peaks with prominent fountain score peak.

## Materials and methods

### Experimental model and subject details

Wild-type fish of AB/TL and mutant strains were raised, maintained, and crossed under standard conditions as described by Westerfield ^73^. Embryos were obtained by natural crossing (4 males and 4 females in 1,7 l breeding tanks, Techniplast). Wild-type and mutant embryos from natural crosses were collected in parallel in 10-15 minute intervals and raised in egg water at 28.5°C until the desired stage. The staging was performed following the Kimmel staging series ^32^. Stages of the mutant embryos were indirectly determined by observation of wild-type embryos born at the same time and incubated under identical conditions. All experiments were performed in accordance with German Animal Protection Law (TierSchG) and European Convention on the Protection of Vertebrate Animals Used for Experimental and Other Scientific Purposes (Strasburg, 1986 ^74^). The generation of double and triple mutants was approved by the Ethics Committee for Animal Research of the Koltzov Institute of Developmental Biology RAS, protocol 26 from 14.02.2019.

### Generation of MZ*sn*, MZ*pn* double mutant and MZ*triple* triple mutant embryos and maintenance of the mutant fish lines

Maternal-Zygotic (MZ) homozygous mutant embryos MZ*sn* (MZ*sox19b* ^m1434^*nanog* ^m1435)^ were obtained in three subsequent crossings. First, MZ*nanog* ^m1435 42^ homozygous null-mutant males were crossed with MZ*sox19b* ^m1434 40^ homozygous null-mutant females. The double heterozygous fish were raised to sexual maturity and incrossed. The progeny developed into phenotypically normal and fertile adults. Genomic DNA from tail fin biopsies was isolated and used for genotyping. We first selected *sox19b-/-* homozygous mutants by PCR-with Sox19b-f1/Sox19br1 primers followed by restriction digest with BbsI, as described in ^40^. To select the double homozygous fish, we used the genomic DNA from *sox19b-/-* homozygous mutants by PCR with Nanog-f2/Nanog-r2 primers followed by restriction digest with NdeI, as described in ^42^. MZ*sn* embryos for experiments were obtained from incrosses of double homozygous *sox19b*-/-; *nanog-*/- fish. The line was maintained by crossing *sox19b*-/-, *nanog* -/- males with *sox19b*-/-, *nanog* +/- females.

MZ*sn* homozygous mutant embryos were obtained in three subsequent crossings. First, MZ*nanog* ^m1435^ homozygous males ^42^ were crossed with MZ*spg* ^m793 41^ homozygous females. *Spg*^m793^ allele carries an A->G point mutation in the splice acceptor site of the first intron of Pou5f3 gene, which results in the frameshift starting at the beginning of the second exon, before the DNA-binding domain. Spg^m793^ is considered to be null allele. The double heterozygous fish were raised to sexual maturity and incrossed. To bypass the early requirement for Pou5f3 in the *spg^m7^*^93^ homozygous mutants, one-cell stage embryos were microinjected with 50-100 pg synthetic Pou5f3 mRNA as previously described ^41^. The fish were raised to sexual maturity (3 months), and genomic DNA from tail fin biopsies was isolated and used for genotyping. We first selected *nanog*-/- homozygous mutants, by PCR with Nanog-f2/Nanog-r2 primers followed by restriction digest with NdeI ^42^. To select the double homozygous fish, we used the genomic DNA from *nanog*-/- homozygous mutants to PCR-amplify the region flanking the *spg*^m793^ allele. We used the following PCR primers: spg-f1 5’-’ GTCGTCTGACTGAACATTTTGC -3’ and spg-r1 5’-’ GCAGTGATTCTGAGGAAGAGGT -3’. Sanger sequencing of the PCR products was performed using a commercial service (Sigma). The sequencing traces were examined and the fish carrying A to G mutation were selected. MZ*pn* embryos for experiments were obtained from incrosses of double homozygous *spg*-/-;*nanog*-/- fish. The line was maintained by crossing *spg*-/-, *nanog* -/- males with *spg-*/-, *nanog* +/- females and microinjecting Pou5f3 mRNA in each generation.

To obtain the triple Maternal-Zygotic homozygous mutant embryos MZ*triple*, MZ*sn* double homozygous males were crossed with MZ*ps* (MZ*sox19b^m1434^spg^m793^* double homozygous females ^40^. The *sox19b-/-*; *spg+/-*; *nanog +/-* progeny was raised to sexual maturity and incrossed. The progeny was microinjected with 50-100 pg synthetic Pou5f3 mRNA at one cell stage, raised to sexual maturity and genotyped as described above. MZ*triple* embryos for experiments were obtained from incrosses of triple homozygous fish. The line was maintained by crossing *sox19b*-/-;*spg*-/-, *nanog* -/- males with *sox19b*-/-;*spg*-/-, *nanog* +/- females and microinjecting Pou5f3 mRNA in each generation.

### Genomic DNA isolation and PCR for genotyping

Genomic DNA was isolated from individual tail fin biopsies of 3 months old fish. Tail fin biopsies or embryos were lysed in 50µl lysis buffer (10 mM Tris pH 8, 50 mM KCl, 0.3% Tween20, 0.3% NP-40, 1mM EDTA) and incubated at 98°C for 10 min. After cooling down Proteinase K solution (20 mg/ml, A3830, AppliChem) was added and incubated overnight at 55°C. The Proteinase K was destroyed by heating up to 98°C for 10 min. The tail fin biopsies material was diluted 20x with sterile water. 2µl of was used as a template for PCR. PCR was performed in 25-50 µl volume, using MyTag polymerase (Bioline GmbH, Germany) according to the manufacturer’s instructions, with 30-35 amplification cycles.

### Zebrafish sperm collection

Adult male fish were anesthetized with Tricain (4% 3-Aminobenzoic acid ethyl ester, pH 6.7) and then positioned with the anal area above. The sperm was taken using a capillary, mixed with 5 µl E400 buffer (9.7 g KCl, 2.92 g NaCl, 0.29 g CaCl_2_ -2H_2_O, 0.25 g MgSO_4_ -7H_2_O, 1.8 g D-(+)-Glucose, 7.15 g HEPES in 1L dH_2_O, pH 7.9) and then with 150µl SS300 buffer (0.37 g KCl, 8.2 g NaCl, 0.15 g CaCl_2_ -2H_2_O, 0.25 g MgSO_4_ -7H_2_O, 1.8 g D-(+)-Glucose 20ml 1M Tris-Cl, pH 8.0, in 1L dH_2_O). The collected sperm was used for nuclei isolation.

### Hi-C library preparation

The embryos were obtained from natural crossings in mass-crossing cages (4 males + 4 females). 5-10 cages were set up per genotype, and the eggs from different cages were pooled. The freshly laid eggs were collected in 10-15 min intervals. Embryos were incubated at 28.5°C and dechorionated with pronase E (0.3 mg/ml) shortly before the desired stage. 400-600 embryos were homogenized in 2 ml 0.5 % Danieau’s with protease inhibitor cocktail (PIC, Roche) and 1 % (v/v) Methanol-free Formaldehyde (Pierce) and fixed for 10 min on a rotating platform. The fixation was stopped with 0.125 M Glycine by shaking for 5 min on a rotating platform. Cells were pelleted on the tabletop centrifuge for 5 min, 500 g, and washed three times with PBST (16mM Na_2_HPO_4_, 4mM NaH_2_PO_4_,0.08% NaCl(w/v), 0.002% KCl (w/v), 0.1% Tween 20, pH 7.5), with protease inhibitors. The cells were lysed for 1 min in 1 ml lysis buffer (10 mM Tris-HCl (pH 7.5), 10 mM NaCl, 0.5 % NP-40) on ice. The pellet was washed 2 times with 1 ml ice-cold 1x PBST. In order to count the obtained nuclei, the pellet was resolved in 1 ml ice-cold 1x PBST, of which 10 μl were diluted 1:1 with 12 μM Sytox® green. The nuclei were scored under a fluorescence microscope using the Neubauer counting chamber. The residual nuclei were snap-frozen in liquid nitrogen and stored at -80 °C. 2.5-3 million nuclei were used for one Hi-C experiment according to the published protocol ^75^.

### Chromatin accessibility changes on fountains in MZ*triple*, MZ*spg* and MZ*nanog* mutants compared to the wild-type

To evaluate the changes in chromatin accessibility on fountains in the absence of Pou5f3, Nanog, or all three zygotic genome activators, we analyzed ATAC-seq data in the respective mutants and the wild-type ^43^ using the European galaxy server ^76^ (ED Fig. 4).

The coverage of ATAC-seq reads at 1460 fountains was scored using MultiCovBed (Bedtools) in four ATAC-seq replicates of the wild-type, three replicates of MZ*spg* at 4.3 hpf (GEO accession number: GSE188364), two replicates of MZ*nanog* and four replicates of MZ*triple* at 4.3 hpf (GEO accession number: GSE215956). Deseq2 ^77^ was used to normalize the reads and compare chromatin accessibility in each mutant to the wild-type. Each fountain was scored as weakened in the mutant if the log2 fold change to the wild-type was negative with FDR < 5%. As a result of this analysis, 1460 fountains were divided into five groups, as shown in ED Fig. 4a:

I. P - downregulated in MZ*spg*,
II. N - downregulated in MZ*nanog*,
III. T - downregulated in MZ*triple*, not in MZ*spg,* and not in MZ*nanog*,
IV. All - downregulated in MZ*triple*, MZ*spg,* and MZ*nanog*,
V. the rest - not downregulated in three mutants.

Group assignments, Deseq2-normalised ATAC-seq values per replicate, average ATAC-seq values, and the values of log2 fold change in each of the mutants compared to the wild-type are listed in the Suppl. Dataset 2 for all fountains and were used for ED Fig. 4d-g.

### GREAT analysis

For predicting the function of cis-regulatory regions within fountains, we used GREAT (Genomic Regions Enrichment of Annotations Tool ^44^ on all ATAC-seq peaks ^43^ overlapping fountains. ATAC-seq peaks were associated with genes using single nearest gene: 50000 bp max extension rule.

### Replication timing data analysis

Replication timing data were from ^38^. Genomic .txt files containing coordinate in danrer 10 (chromosome number in column 1 and chromosomal location in column 2), and replication timing value in column 3, were obtained from GEO GSE85713. Timing value represents S-phase copy number relative to G1 sample, normalized to a genome mean of 0 and standard deviation of 1. Files were converted to .bedgraph format and danrer 11 assembly using liftover utility ^78^. Genomic coordinates of the initiation zones were obtained from Chris Sansam. Heat maps and profiles were built using deep tools ^79^.

### Hi-C data mapping

Hi-C reads were mapped with Open2C *bwa-mem* ^80^ and *pairtools*^81^-based pipeline *distiller-nextflow* version 0.3.3. For mapping, we used UCSC’s danRer11 assembly with removed unplaced contigs ^82^. For parsing, we used the walks policy “all” for parsing pairs ^81^. This increased the library pairs yield by parsing all possible pairs from each sequenced read pair. Cool files were created by *cooler,* and replicates were merged by summation ^83^. Normalization was done by iterative correction ^84^ with default parameters (minimal count per genomic bin 0, minimum number of non-zero pixels per bin 10, maximum median absolute deviation of pixel values per bin 5, both cis- and trans-contacts included, tolerance 1e-05 and 200 maximum iterations).

Processed coolers from this work are available at GEO ^85^ https://www.ncbi.nlm.nih.gov/geo/query/acc.cgi?acc=GSE195609. We also re-processed data from previously published zebrafish embryogenesis works ^4,33^, and made the output available through the Open Science Foundation ^86^ website: https://osf.io/mt4vf/.

### Hi-C data processing

#### Hi-C data visualization

Normalized Hi-C signal was visualized with HiGlass genomic browser ^87^ (Figure 1a,b) and resgen ^88^ (SI Fig. 4) in log-scales and “fall” colormap.

#### Centromere positioning in *Danio rerio*

We visualized Hi-C maps for *Danio rerio* in HiGlass genomic browser ^87^ and manually marked the locations of centromeres following criteria: (i) centromeres must be located far from chromosome ends; (ii) regions around centromeres should have enriched interactions with regions around other centromeres in trans; (iii) centromeres should have unmapped DNA as a sign of centromeric repeats. The beginning and the end of the unmapped DNA served as centromere start and end, respectively. We later used this annotation to plot the Rabl configuration.

#### P(s) curves and derivatives

To calculate P(s) curves, also named scalings, for ED Fig. 1d, we used *cooltools* ^36^ v0.5.4 *expected_cis* at 1 kb resolution. We removed all Hi-C contacts shorter than 2 kb, binned contacts in log space to produce evenly sized points in double log coordinates, then smoothed contacts with Gaussian kernel with sigma 0.05 to even out noisy interactions at the ends of chromosomes and applied aggregation per chromosome arm (defined as in “Centromere positioning”). P(s) derivatives were calculated by taking the gradient at each point of the resulting P(s) plot. For each dataset, the approximate distance ranges of peak locations were manually marked and then validated by searching for local maxima of the derivative plots at distances under 120 kb.

Note that saddle plots (ED Fig. 1b) and snipping Hi-C data (for average pileup and fountain calculation) are separate analyses that also utilize expected P(s) (see below). However, for these analyses, we did not smooth or aggregate contacts.

#### Removal of poorly mapped and surrounding genomic regions

While analyzing centromere positions, we noticed that the Hi-C data frequently had rearrangements and poorly mapped regions, noticeable as sharp shapes and edges in the smooth-looking map. Further analysis, thus, required the exclusion of poorly mapped regions. For that, we created a track of unmappable genomic bins (or “bad bins”).

Firstly, we defined bin as “bad” if it was impossible to normalize through the default cooler balancing (see criteria in the “Hi-C data mapping”) in at least one of the replicate or merged datasets. Secondly, we marked all genomic regions located at least 50 Kb away from these bins as bad because they might be affected by balancing problems due to the absence of short-range interactions. Moreover, bad bins frequently had genomic rearrangements around them, which might affect TAD and fountain calling.

These two criteria resulted in the extended list of 49843 genomic bins subject to removal from most of the Hi-C analyses (out of the total 134526 bins, with 4799 bad bins located solely at chr4).

#### Building developmental trajectories

To build developmental trajectories, we calculated pairwise SCC (stratum-adjusted correlation coefficient) ^89^ between all zebrafish Hi-C datasets from previous studies and this work (including replicates and merged datasets). For that, we used the R library HiCRep ^89^ with a minimum genomic separation of 0, maximum separation of 5 Mb, window size of 3, and data resolution of 100 Kb. The resulting SCC was input to PCA analysis with *sklearn* ^90^. The first two PCs jointly explained around 79% of the variance, with a substantial drop at PC3: PC1 47.7%, PC2: 23.1%, PC3: 17.4%, PC4: 4.1% (Fig. 1f). Note that the PCa was done using all zebrafish Hi-C available experiments to us, including Hi-C of mutants. Mutants were excluded from visualization for simplicity.

#### Average Rabl configuration

To assess Rabl configuration (ED Fig. 1a,c), we plotted average Rabl pileups. We extracted the trans interactions for each pair of chromosome arms at 250 Kb resolution (per-arm snippets). We reshaped per-arm snippets of different sizes to the 200×200 squares with *cooltools* ^36^, which is a common strategy for averaging Hi-C pileups of different sizes ^91^. We then averaged all snippets and displayed log2 over expected (average number of trans-interactions).

The centromeric score measures the enrichment of contacts between centromeres of different chromosomes relative to all trans interactions. To calculate the centromeric score, we started with the average Rabl pileup (reshaped and normalized), took a 40×40 window around its center, and calculated the median value.

#### Compartment calling and saddle plots

For compartment calling (ED Fig. 1b,c), we used *cooltools* cis eigendecomposition with *eigs_cis* ^36^. We used cis decomposition and not trans to avoid the influence of prominent Rabl configuration.

Phasing of the eigenvector is an important step of compartmental calling that defines the sign of the eigenvector with A compartment ^84,92^. Phasing of the mammalian first eigenvector is based on correlating the eigenvector with either number of genes or GC content ^84,92,93^, because both are enriched in the A compartment of mammals ^36^. For example, in mouse, the first eigenvector at the late 2-cell stage of embryo development ^7^ highly correlates with both GC content (Pearson corr. 0.64 at 25 kb bin size) and coverage by genes (Pearson corr. 0.28).

Surprisingly, the first eigenvector of zebrafish Hi-C at 5.3 hpf does not correlate with GC content (Pearson corr. of -0.037 at 25 kb bin size) and poorly correlates with coverage by genes (P. corr. 0.137). However, the first eigenvector at 5.3 hpf correlates well with replication timing at shield stage ^94^ (P. corr. 0.59). Thus, we phased the first eigenvector of Hi-C maps with replication timing instead of conventional GC content. 11 hpf first eigenvector correlated the best with bud RT (P. corr 0.65), and 25 hpf first eigenvector correlated the best with RT measured at 28 hpf (P. corr 0.65) ^94^.

Saddle plots (ED Fig. 1b) were constructed by aggregating observed over expected cis Hi-C signal at 100 kb (calculated by normalizing balanced Hi-C signal by expected with no smoothing), based on grouping into 20 groups by first eigenvector for corresponding stage (see details in ^36^).

Compartment score (ED Fig. 1c) was defined as the mean log2 number of AA and BB interactions in saddle plots minus the mean log2 number of AB and BA interactions. The transition point between A and B compartments was defined by zero of the first eigenvector. For the distribution of compartmental scores, we calculated the compartment scores for each chromosome and plotted the resulting values as distributions.

#### Insulation score

To capture the general properties of the scalings in embryogenesis, we first calculated insulation scores ^66^ with *cooltools* ^36^ at 5 Kb resolutions for each replicate of each developmental stage of zebrafish. We set the window size to 200 Kb. Next, we found local peaks in the insulation score profiles, considered peaks reproducible between replicates as boundaries, and used them for insulation strength characterization in each experiment (ED Fig. 1).

For pileups (Fig. 2a), we first constructed *bigwig* files with insulation scores with *bioframe* ^95^ and then aggregated the signal in the genomic windows by *pybbi* ^96^, as done in ^97^.

#### TAD calling

For the detection of boundaries and TADs in Fig. 2a, SI Fig. 8, we performed more rigorous TAD boundary calling. We processed each replicate of each developmental stage with *cooltools*-based and insulation score-based *HiChew* tool ^98^ at 10 kb, as explained below. We set the expected TAD size parameter of 200 Kb ^33^ and automatically assessed the optimal parameters of TAD calling to match the expectation. Next, we removed the TAD boundaries that overlapped with the bad bins. All code for this analysis is deposited in the GitHub repository: https://github.com/encent/danio-2022.

Function *hichew.calling.boundaries* produces TAD boundaries calling given a certain resolution and the expected TAD size parameter. By iterating over the insulation window size grid, it finds for each chromosome the insulation window size parameter that minimizes the difference between the user-defined expected TAD size parameter and the median TAD sizes obtained while iterating the grid. The resulting TAD boundaries annotation is calculated with *cooltools* functions *calculate_insulation_score* and *find_boundaries* using the insulation window size parameters found for each chromosome, as explained before. All code for TAD boundaries calling using *HiChew* utility is deposited in the Jupyter notebook: https://github.com/encent/danio-2022/blob/main/src/TAD_boundaries_calling.ipynb.

For the confident set of TAD boundaries at 11 hpf in Fig. 2a,b, ED Fig. 2c, we further selected only those boundaries that contain at least one ATAC-seq peak supported by CTCF motif (see “CTCF binding inference from ATAC-seq” section).

#### Fountain calling with *fontanka*

##### Reference fountains

We first arbitrarily picked regions of chromosomes 1 and 2 and manually marked fountains there. In total, we collected 34 fountains from chromosome 1 and 18 fountains from chromosome 2. This fountain set was used as a reference fountain for fountain calling control.

##### Fontanka protocol

We designed *fontanka*, *cooltools*^36^-based tool for fountains calling in Hi-C maps (Figure 2b). Although *fontanka* is designed to call any on-diagonal pattern in a Hi-C map, we applied it here to call fountains. Fontanka has four principal steps (ED Fig. 1e):

(i) **Snippets extraction.** *Fontanka* rolls a square window of specified size along the main diagonal of the Hi-C matrix and extracts observed over the expected signal. Each window is assigned to the genomic point of its center.
(ii) **Convolution.** Instead of insulation diamond window implemented in *cooltools* insulation ^36^, *fontanka* performs two types of convolution:

(a) Fountain score calculation. Each snippet is convolved with the **fountain mask**, which produces a **fountain score** for the corresponding genomic region. In this work, we used average pileup for all reference fountains as a fountain mask. The convolution score can be based on the summed product of the snippet with a mask, mean square error, and Pearson or Spearman correlation. Here, we used Pearson correlation as a measure for fountain score, which proved to be the best in preliminary tests of the library. Thus, the fountain score can have values between (-1, 1), where 1 is the best correlation with the fountain pattern, and 0 is the absence of correlation.
(b) Noise score calculation. Since Hi-C maps frequently displayed rearrangements relative to the default danRer11 genome, we aimed to filter out potential rearrangements as a noise source in our data. For that, we applied another convolution round with the Scharr operator, approximating the gradient of the Hi-C map ^99 100^. We averaged the Scharr operator by two orientations (vertical and horizontal) for each window pixel, resulting in a **Scharr score**. The Scharr score measures the noise and sharpness of the signal in the corresponding window. Too noisy or sharp patterns in Hi-C snippets result in high noise scores, while smooth Hi-C maps will have Scharr scores closer to zero.

As a result, each fountain candidate thus has two characteristics: fountain score and noise score (ED Fig. 1f). The higher the fountain score, the more pronounced the fountain-like structure. The smaller the Scharr score, the less noisy the local Hi-C map and the less probable the rearrangements at the corresponding genomic locus.

(iii) **Peak calling.** *Fontanka* detects local maxima in the fountain score by *cooltools find_peak_prominence* function ^36^. This function detects peaks in the 1D genomic signal of the fountain score and assigns **peak prominence** to each one. The resulting peaks were considered candidate fountains (ED Fig. 1e).
(iv) **Thresholding and filtration.** Candidate fountains contain potentially very weak and noisy false calls. In order to minimize the contribution of false calls, we set strict requirements for fountains and select the most confident set of fountains (ED Fig. 1e).

(a) We first remove the fountains with weak prominence. We noticed that the distribution of the fountain peak prominence is a mixture of two distributions: small noisy calls and long tail of high values of potential true peak calls. To separate those distributions, we applied Li’s iterative method for finding the separation point by using the slope of the cross-entropy ^101^, as implemented in *scikit-learn* ^102^.
(b) Next, we remove fountains that fall too close (<50 Kb) to the nearest bad bin (see “Removal of poorly mapped and surrounding genomic regions” section of Methods).
(c) We remove all the fountains that have a negative correlation with the fountain mask.
(d) We next require that the fountains detected in the merged dataset is detected in individual replicates with an offset of 20 kb (on either side from the fountain).
(e) We apply a filter for potential genomic rearrangements and misassembly problems. Genomic rearrangements should be present at all the stages, including 2.75 hpf that does not have visible non-artifactual on-diagonal patterns in Hi-C. Thus, any fountain detected at 2.75 hpf is probably a result of the erroneous call of genomic rearrangement as a fountain. We removed those fountains at 5.3 hpf that fall within 20 kb from the nearest constant “fountain” at 2.75 hpf (reproducible between replicates at this time point).
(f) Another filter for potential genomic misassembly problems filters out the top 25% of candidate fountains based on their Scharr score.

Out of 3391 candidate fountains at 5.3 hpf passing filters iv.a-iv.c, 2199 were confirmed in replicates (64.8%, filter iv.d). 1110 genomic bins were found as constant “fountains” at 2.75 hpf. Filter (iv.e) resulted in 1947 candidates, out of which 1460 were finally selected for the analysis (iv.f). The final list of fountains with fountain scores and fountain peak scores is provided as Supplementary Dataset 1.

15 out of 34 reference fountains at chr1 (44.1%), and 10 out of 18 reference fountains at chr2 (55.5%) were successfully found by *fontanka* with all filtration steps (Extended Data Fig. 1b).

Since Open2C software are under ongoing development, we fixed the software versions to fontanka v0.1, bioframe v0.4.1 ^95^, cooler v0.8.11 ^83^, and cooltools v0.5.2 ^36^ in this work.

##### Comparison of fontanka and Chromosight

We noted that the fountain structures are similar to hairpins ^103^, and the hairpin calling algorithm (*Chromosight*) can be applied to call fountains in our dataset. With *Chromosight* v1.6.3 and default parameters, we identified 9 out of 34 reference fountains at chr1 (26.5%) and 7 out of 18 reference fountains at chr2 (38.9%). Notably, *Chromosight* output had an average score of 0.14 and an average pileup that did not resemble the average reference fountain. *Chromosight* results are posted online at https://github.com/agalitsyna/fontanka/blob/master/examples/01_fontanka_vs_chromosight.ipynb.

##### Comparison of fontanka and protractor tool

Another algorithm for fountain-like structure calling was proposed for jets ^29^. It relies on calculating the contacts at certain distances away from the base (protractor tool). To replicate this approach, we calculated the number of Hi-C interactions normalized by expected in a line perpendicular to the main diagonal of the Hi-C map for the whole genome. Then, we compared the fountain average protractor versus all non-fountain bins. Indeed, the protractor tool showed enrichment of the contacts at distances up to 150 kb (SI Fig. 7). We summed up the protractor tool average over all distances up to 200 kb and correlated it with the fountain score (ED Fig. 1g). Both scores correlated with 0.32 Pearson correlation, 0.44 Spearman correlation.

#### Enrichment of developmental regulatory elements at fountains

For the analysis of regulatory elements at the fountains (Fig. 2c, ED Fig. 2e, Fig. 4g-h), we started with developmental regulatory elements from ^39^, also called predicted ATAC-seq-supported developmental regulatory elements (PADREs). Briefly, PADREs are ATAC-Seq at the dome stage, annotated by ten ChromHMM states. ChromHMM states were obtained from the set of histone modifications measured in *Danio rerio* development.

For the enrichment analysis, we first split the genome into non-overlapping 10 kb bins. Then, we calculate the coverage of each genomic bin by the regulatory element of a certain type with the *bioframe coverage* tool ^95^. Then, for each bin, we calculate the relative **enrichment of the regulatory element** (the ratio between coverage of the bin by a regulatory element divided by coverage by open chromatin of any type). We then average the enrichments for all bins at fountain bases.

As a control, we randomize the positions of fountains and calculate the average enrichments again for each state. This procedure preserves the distribution of ATAC-Seq peak sizes and the relative contribution of each state to PADRE annotation. For FDR calculation, we approximated the control distribution by Gaussian and calculated the probability of obtaining an observed value or larger.

#### Hi-C snipping and average pileup

To snip the Hi-C maps (across the whole manuscript), we used *cooltools pileup* function ^36^. Pileup takes one or multiple genomic locations and returns a square window of specified size around these locations.

To build average pileups over genomic loci (e.g., fountains), we took the mean of corresponding pixels of all windows snipped around fountains.

#### Differential fountains in MZ*triple*

To define the fountains that go down, stay unchanged, or go up in MZ*triple* (ED Fig. 3e), we first calculated the fountain scores by fontanka (as described above). Then, we calculated the difference in fountain scores between WT and MZ*triple* for each genomic bin. We then plot the distribution of all differences and fit it using the Gaussian distribution (by *scipy stats norm* ^104^ function). We then defined ’DOWN’ fountains as those that have a drop in fountain score more than 1.5 standard deviations from zero and ’UP’ as those that have a growth of fountain score more than 1.5 standard deviations from zero. The rest of the fountains were considered unchanged.

### ChIP-Seq data analysis

We re-analyzed ChIP-Seq data from ^4,22,25,27,67^ with *chipseq-nf* 1.2.1 with default parameters ^105^. The processing mode was set either to single-end or paired-end based on raw sequencing annotation in the SRA database (see Suppl. Dataset 6).

### CTCF binding inference from ATAC-seq

Unfortunately, at the time of this research, CTCF ChIP-Seq data is not available for most stages of development of zebrafish that we study (except one dataset for 10 hpf from ^27^). However, multiple ATAC-Seq profiles are available from the Danio Code database ^106^. This allowed us to infer CTCF binding from ATAC-Seq data by assuming that the ATAC-Seq peak with the strong CTCF motif has CTCF bound at this stage.

First, we called CTCF motifs in the danRer11 genome. Generic vertebrates CTCF motif (JASPAR MA0139.1) instances were called by JASPAR whole-genome *PWMScan*-based ^107^ motif scanner ^108^, as described in ^109^. The background GC content was set to 0.317 (A and T frequencies) and 0.183 (G and C). We used a p-value threshold of 5e-02 and an r threshold of 0.8.

Next, we downloaded ATAC-Seq peaks from DCC database ^106^ in narrow peak format for 4.5 hpf (DCD019127DT) and 12 hpf (DCD019077DT). Each ATAC-Seq peak was marked as CTCF-bound if it had at least one CTCF motif.

The selection of parameters for motif calling was made by benchmarking against CTCF ChIP-Seq peaks obtained at 10 hpf from ^27^ (re-processed with *chipseq-nf* with default parameters ^110^).

For coverage pileups in ED Fig. 2c, we stored the CTCF-containing peaks with their abundance as bigwig files with *bioframe* ^95^ and then aggregated the signal in the genomic windows by *pybbi* ^96^.

### Simulations of loop extrusion

Simulations of loop extrusion (Fig. 5, ED Fig. 5-6) were based on the *OpenMM*-based ^111^ Python library *Polychrom* ^112^. We split it into five steps: (1) 1D simulation of extruders movement on the genome, (2) 3D simulation of polymer, (3) *in silico* reconstruction of interaction probabilities, (4) assessment of goodness of fit of the simulations to real data, (5) parameter sweep of the model.

The implementation is available at https://github.com/agalitsyna/polychrom_workbench under the *targeted_extrusion* directory.

#### 1D simulation

We simulated the translocation of extruders on a 1D lattice ^113^, where each point represented a genomic position. Each position can either (1) have a regular role or (2) serve as a loading platform for the targeted extruder. For regular positions, we assumed uniform probabilities of loading and unloading. For loading platforms, we set up the probability of cohesin landing increased by enrichment factor.

To simplify the model, we assumed (1) immediate reloading of unloaded cohesin, and (2) separate pools of extruders that can load at regular positions versus loading platforms. Before running the simulations, we ensured that cohesins were fully loaded on a lattice.

As in our previous work ^113^, simulations were organized in rounds, with all cohesins performing translation with both legs in opposite directions at each round. Each simulation lasted for 10 010 000 translocation rounds, and we sampled each 10th step as a readout of 1D simulation. We removed the first 10000 steps to ensure that we sample the steady state of loop formation and not the earlier stages of loop formation and maturation ^114^.

The 1D lattice size was 50000 monomers, with each monomer corresponding to 1 Kb of DNA (thus, the whole simulated molecule was 50 Mb).

The size of the loading platform was set to 1 monomer (1 kb), and the platforms were distributed each 500 kb to correspond to the mean distance between fountains in Hi-C data. Technically, we generated 20 groups of the size 2500 to allow averaging over multiple groups in a single conformation when calculating the average fountain, as we proposed previously ^12^.

For the basic simulations of targeted loading, we assumed that there are (1) no boundaries that stall cohesin legs, (2) only other cohesins can stall the legs of cohesins, (3) legs of cohesin (if not stalled) are moving simultaneously, (4) free leg of cohesin with a stalled leg can move.

For simulations with random barriers, we assumed that (1) cohesins load only at the loading platforms, (2) random barriers are placed at each genomic position in a 100 kb window around the platform, (3) random barriers stall the cohesin leg with some probability (best fit to real data obtained for the stalling probability of 0.005, see Fig. 5c, *middle*).

For simulations with decoupled legs of cohesin, we assumed that (1) cohesins load only at the loading platforms, (2) the probability of stepping is set for each leg independently (best fit to real data obtained for the probability of stepping set to 0.99, see Fig. 5c, *right*). We note, however, that by decreasing the probability of stepping, we also decreased the speed of extruders, which may explain why we obtained the stepping probability close to 1.

#### 3D simulation

As in our previous works ^12,37,114^, we represented chromatin as a polymer with spherical monomers connected by harmonic bonds with stiffness and soft-core repulsive potential. Simulations were performed with variable Langevin integrator in the periodic boundary conditions ^12^ with volume density 0.1. For simulations and setting up the bonds, we used the OpenMM-based ^111^ Python library Polychrom ^112^. A harmonic bond connected the two monomers held by the extruder. The number of 3D-simulation time steps per 1D simulation step was set to 200 ^46^.

#### *In silico* reconstruction of interaction probabilities

To calculate contact maps, we recorded all the pairs of monomers located closer than 5 monomer radii. We then averaged the interactions over multiple equivalent groups to obtain a simulated 2.5 Mb region with fountains. To obtain a simulated average fountain at 10 kb resolution, we snipped 200 kb windows around the loading platforms and coarse-grained them by a factor of 10. We then normalized the snippets by expected.

#### Goodness of fit of the simulations to real data

The goodness of fit was assessed by the correlation of the simulated average fountain with the Hi-C average fountain. For that, we compared each *in silico* Hi-C of the simulated fountain with an average reference fountain. As a quality measure, we calculated (1) the Spearman correlation between matrices of average fountains, (2) protractor sections of average fountains, and (3) cross-sections of fountains at characteristic distances from diagonal (Sketch: ED Fig. 5c, results: Fig. 5d-e, ED Fig. 5b-c, more illustrative examples for different simulation parameters: ED Fig. 6). Both matrices were taken at 10 kb resolution, and *in silico* Hi-C matrices were coarse-grained for this purpose (as opposed to non-coarse-grained visualizations of average simulated pileups, which are displayed at 1 Kb resolution).

For additional validation, we checked the interactions between loading platforms (fountain bases) by snipping off-diagonal in silico Hi-C matrices and verified that there was no enrichment of interactions (ED Fig. 5d).

#### Parameter sweep

The simulations were organized as a set of Python scripts: 1D extrusion, 3D simulations, and 2D contact map construction. For the 1D extrusion, we could vary lifetime, separation, and the level of targeted loading. For 3D simulations and 2D contact map construction, we fixed the parameters. To automate the parameters sweep, we implemented a snakemake ^115^ pipeline combining all three steps. Tested parameters and the resulting goodness of fit are presented in Fig. 5d (the color represents the mean square error (MSE) of the protractor) and ED Fig. 5b (the color represents the Spearman correlation coefficient).

