## Supplementary Information for "Extrusion fountains are hallmarks of chromosome organization emerging upon zygotic genome activation"

for

Aleksandra Galitsyna #1

1 - Institute for Medical Engineering and Science, Massachusetts Institute of Technology, Cambridge, MA 02139

### -

### I. Fountain-like structures in other biological systems

We define fountains as *patterns of contacts that emanate from a single genomic locus and broaden with distance from the diagonal.* The fountain is a novel pattern of chromatin organization, which is as widespread in zebrafish embryogenesis as TADs, stripes, and dots are in, for example, differentiated mammalian cells [1,2]. We acknowledge that fountains and fountain-like structures might be present in other biological systems, and the mechanisms of their formation might be different from the targeted cohesin extrusion that we propose here.

On the whole-chromosome scale, *Rabl configuration* of some eukaryotes [3,4] and *hairpins* in bacteria [5–7] resemble fountains. Both patterns represent the arms of chromosomes aligned with each other.

Rabl configuration is a consequence of memory of the mitotic organization, where chromosomes in interphase keep the alignment similar to that in the mitotic spindle. That mechanism, supposedly, does not require tethering of the arms or targeted loading of the extruder [3]. Although Rabl configuration is prominent in zebrafish (as we and others [8] report), it is unrelated to the relatively small (<200 Kb in length) fountains at 5.3 hpf.

The alignment of chromosomal arms in bacteria, or *hairpin*, results from the loading of SMC complexes by ParB protein at a centromeric locus with parS sites. This system was thoroughly explored through simulations of targeted loop extrusion [6]. However, these findings cannot be transferred to zebrafish, where multiple much smaller fountains are scattered throughout multiple chromosomes (Fig. SI 1a).

Other previously reported fountain-like structures at smaller scales are *local interaction patterns (LIPs)* [9] , *flares* [8], *plumes* [10], and *jets* [11] (see Table SI 1).

Induction of double-stranded breaks (DSBs) in *S. cerevisiae* leads to the formation of *local interaction pattern (LIP)*, enriched interactions emanating from the DSB. LIPs are 25-Kb in size, and depend on homologous recombination factors, but not cohesin (although cohesin accumulates at their bases). The mechanism of LIPs formation might be, thus, unrelated to active extrusion.

The chromatin of zebrafish sperm cells is folded into at least 333 *flares* [8]. These structures are specific to sperm cells and are absent at the same positions in developing embryos [8]. Visual inspection of the reported Hi-C maps suggests that flares in sperm cells span from hundreds of thousands to millions of nucleotides. These scales are still much larger than we observe for the fountains here. Interestingly, although H3K27ac was enriched at flares bases, Smc3 was not [8]. Thus, the formation mechanism might differ from the targeted extrusion proposed here.

Upon degradation of both Wapl and CTCF in mouse embryonic stem cells (mESCs), the regions with high openness form *plumes* [10] (bioRxiv preprint). Like fountains in zebrafish, plumes are transient (appear at 6 hours after induction of degradation and disappear at 96 hours). The bases of plumes are enriched in Rad21 in untreated mESCs and lose this enrichment in Wapl/CTCF degradation [10]. Moreover, plumes disappear upon Rad21 degradation [10], suggesting they are formed by active extrusion by cohesin.

Primary mouse DP thymocytes have at least 38 *jets* of 1-2 Mb in size in wild-type cells [11]. Jets depend on Rad21 and become longer upon CTCF knockout [11]. Simulations of targeted loop extrusion from Guo et al. [11] qualitatively explain the jets' formation. For example, jets get more dispersed with the distance from the diagonal, suggesting that desynchronization of cohesin arms is required for their proper shaping, as we find for fountains in this work. The simulations in Guo et al. [11] assume the desynchronization of cohesin arms as an intrinsic property of cohesin. However, they do not explore alternative scenarios for the shaping of the disperse form of fountains/jets.

Critically, Guo et al. [11] do not define their extrusion dynamics and length scales in physical terms. For example, the loading platform for cohesin is set to either 100 (“narrow” platform) or 1000 (“broad” platform) monomers (arbitrary units and not genomic coordinates). From the visual comparison of simulated and Hi-C maps in this work, one could assume that one monomer corresponds to 800 bp (the average window analysis for jets is 4 mb for Hi-C versus 5000 monomers for simulations). Thus, the “narrow” platform in these simulations is, in fact, 80 kb, which is much larger than what we report for fountains in zebrafish (10 kb platform, single monomer). Finally, in contrast to Guo et al. [11], our fitting simulation parameters to the Hi-C data allow us to estimate a cohesin processivity of 150 kb at 1 kb/s speed and enrichment of targeted loading by a factor of 10.

Finally, consistent with the cohesin model of fountains formation, Isiaka et al. [12] observe 1263 fountains at active enhancers of nematode *C. elegans*, and demonstrate that fountains are cohesin-enriched and depend on cohesin.

| **System and condition of fountain-like structures appearance** | **Fountain basis** | **# of fountains per genome** | **# of fountains per 1 Mb of genome (if relevant)** | **Reference** |
|---|---|---|---|---|
| DNA double-strand break formation in *S. cerevisiae* | Double-strand breakpoint | 1 | - | Piazza et al., 2022 [9] |
| Zebrafish sperm cells | H3K27ac-enriched regions | 333 | ~0.26 | Wike et al., 2021 [8] |
| Mouse cells with Wapl + CTCF degron | Open chromatin regions (OCIs, defined as high-ATAC-seq-region flanked by low-ATAC-Seq) | At least several of them; detectable as an average of 105 OCIs | ~0.04 | Liu et al., 2021 [10] (bioRxiv preprint) |
| Small DP thymocytes | Open chromatin regions, also enriched in H3K27ac | 38 | ~0.01 | Guo et al., 2022 [11] |
| Nematode *C. elegans* | Active enhancers, enriched in cohesin | 1263 | ~13 | Isiaka et al., 2023 [12] (bioRxiv preprint) |
| Zebrafish embryos upon ZGA | Open chromatin regions enriched in H3K27ac, H3K4me1 and pioneering factors binding | 1460 | ~1.13 | This work |

**Table SI 1.** Fountain-like structures reported in the literature. Genome size was a sum of all chromosome lengths (as reported by UCSC Genome Browser [13]), excluding mitochondrial chromosomes.

Some works report fountain-like structures at the average pileups, which might also represent fountain formation in chromatin. For example, Murine Endogenous Retroviral Elements with a leucine tRNA primer binding site (MERVL) produce an average Hi-C fountain in embryogenesis (late 2-cell and 8-cell, bioRxiv preprint [14]). Interestingly, Micro-C also reveals a fountain-like average at the binding sites of Nanog and, to a lesser extent, Med12 in mESC [15]. However, in both these works, the fountain-like pattern emerges as an average and might be a consequence of averaging centers of TADs, as we show in Fig. SI 1-3.

Interestingly, Micro-C exposes the features of the average fountain at the resolution of 200 bp, which is much below the resolution possible in Hi-C analysis (10 kb in this work) [15]. At nucleosome resolution, Nanog in mESC has enriched interactions from ~10 kb to ~100 kb, but now at shorter distances. Thus, fountains might have an intricate inner structure. We anticipate that further studies of individual fountains with Micro-C will shed new light on the mechanism of their formation.

Finally, fountain-like structures have also been reported as an outcome of the biophysical modeling of chromatin.

Goychuk et al. [16] model sequence-dependent correlated active forces in chromatin and observe fountain-like structures concomitant with compartments formation. Fountain-like structures originate from small active regions by spontaneous loops by a local hot spot of activity.

Brahmachari et al. [17] observe fountain-like structures in the models with temporally correlated active forces. Fountains are formed at the elements with correlated motion, and become stronger with increased correlation.

Although both these models do not explicitly include loop extrusion (a non-equilibrium process [18]), they might represent the same class of models with non-equilibrium mechanisms. Further work is needed to establish whether fountain-like structures in chromatin are a common signature of non-equilibrium processes in chromatin.

#### II. Emergence of a fountain as an average pileup

An *average pileup* is a commonly used instrument for Hi-C/Micro-C and other capture techniques analysis [19–22]. Average pileup relies on averaging fragments of Hi-C maps (*snippets*) between different genomic locations. This tool has been widely applied in comparative studies between different treatments and conditions [23] or between different genomic locations within a single Hi-C/Micro-C map [15]. However, the resulting average is not representative of the conformation of a single genomic locus. It is important to avoid potential misinterpretation of average pileups.

Fountains are the predominant features of Hi-C maps after ZGA (5.3 hpf) of zebrafish embryos, enabling us to perform genome-wide fountain calling with *fontanka*. With *fontanka*, we call individual genomic locations bearing the fountain signature in the surrounding ±200 kb of the Hi-C map. Visual inspection confirms that snippets have fountains, except for several false positives (see Fig. SI1a of this Supplementary). As expected, when averaged, individual fountains produce the average fountain structure with a large fountain score (Fig. SI1b).

However, the average fountain does not guarantee the presence of fountains at chromosomes. Individual genomic loci that look nothing like fountains may produce the appearance of an average fountain-looking structure at the average pileup. For example, centers of TADs at 11 hpf average to the fountain-looking structure on a pileup (Fig. SI2). This property arises because (1) the centers of TADs have enrichment of contacts between the surrounding regions, and (2) TAD boundaries are located at random distances, washing off the average enrichment nearby. While individual snippets of the pileup do not resemble fountains (Fig. SI2a), an average pileup looks like a fountain (Fig. SI2b). This effect is visible only when a large number of genomic positions with TADs of different sizes is averaged but can be avoided if: (1) individual snippets are inspected visually or (2) snippets with similar TAD sizes are averaged (Fig. SI3a-d).

Thus, the presence of the fountains in Hi-C maps should be confirmed at the level of individual genomic loci and not only by the appearance of the average pileup.

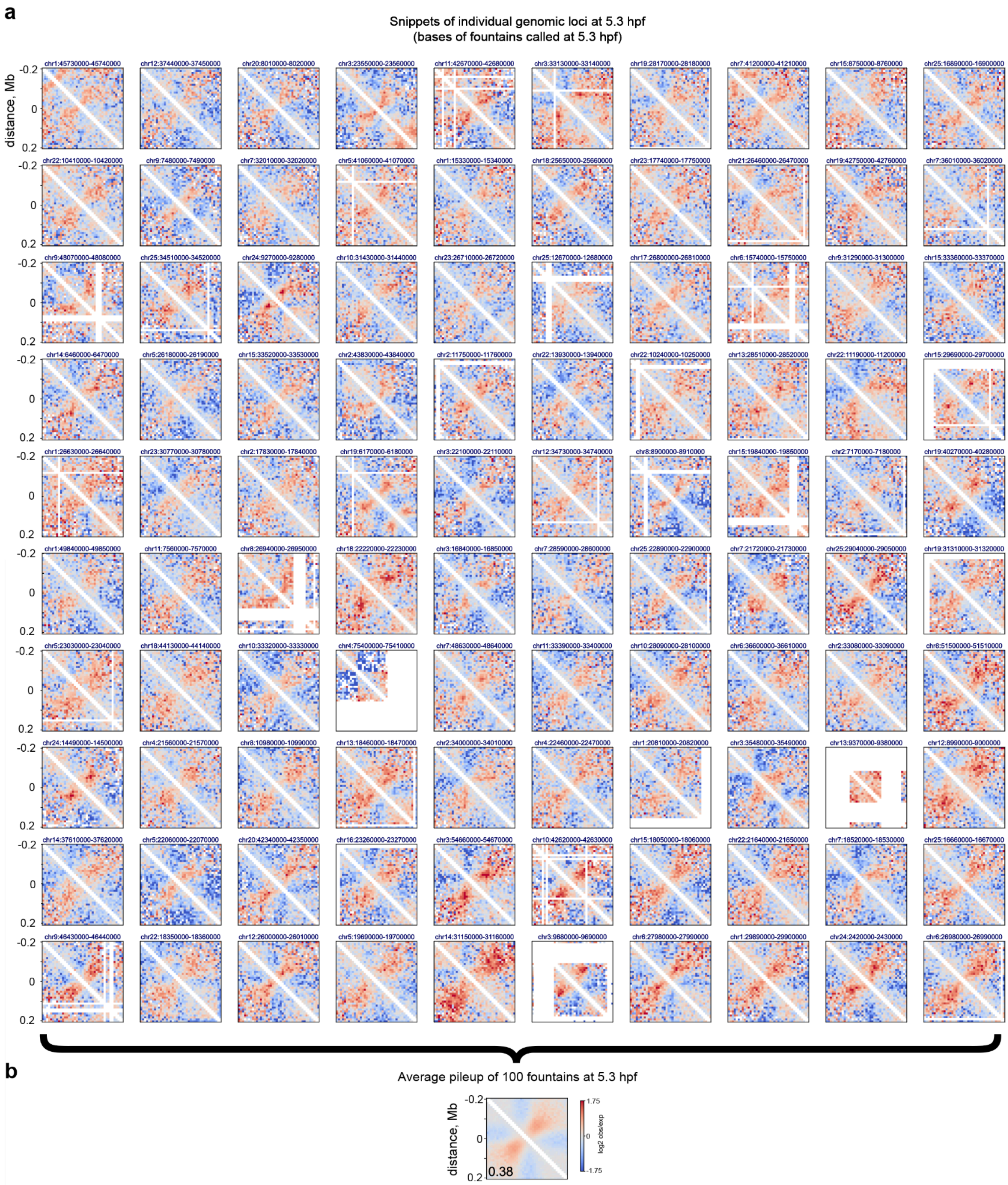

**Figure SI 1. Averaging individual fountains at 5.3 hpf.**

a. Snippets of Hi-C maps of 100 fountain bases called at 5.3 hpf. Each snippet is named by the genomic position of the fountain base.

b. Average pileup with average fountain. The average resembles individual conformations. The number in the corner represents the strength of the fountain signature.

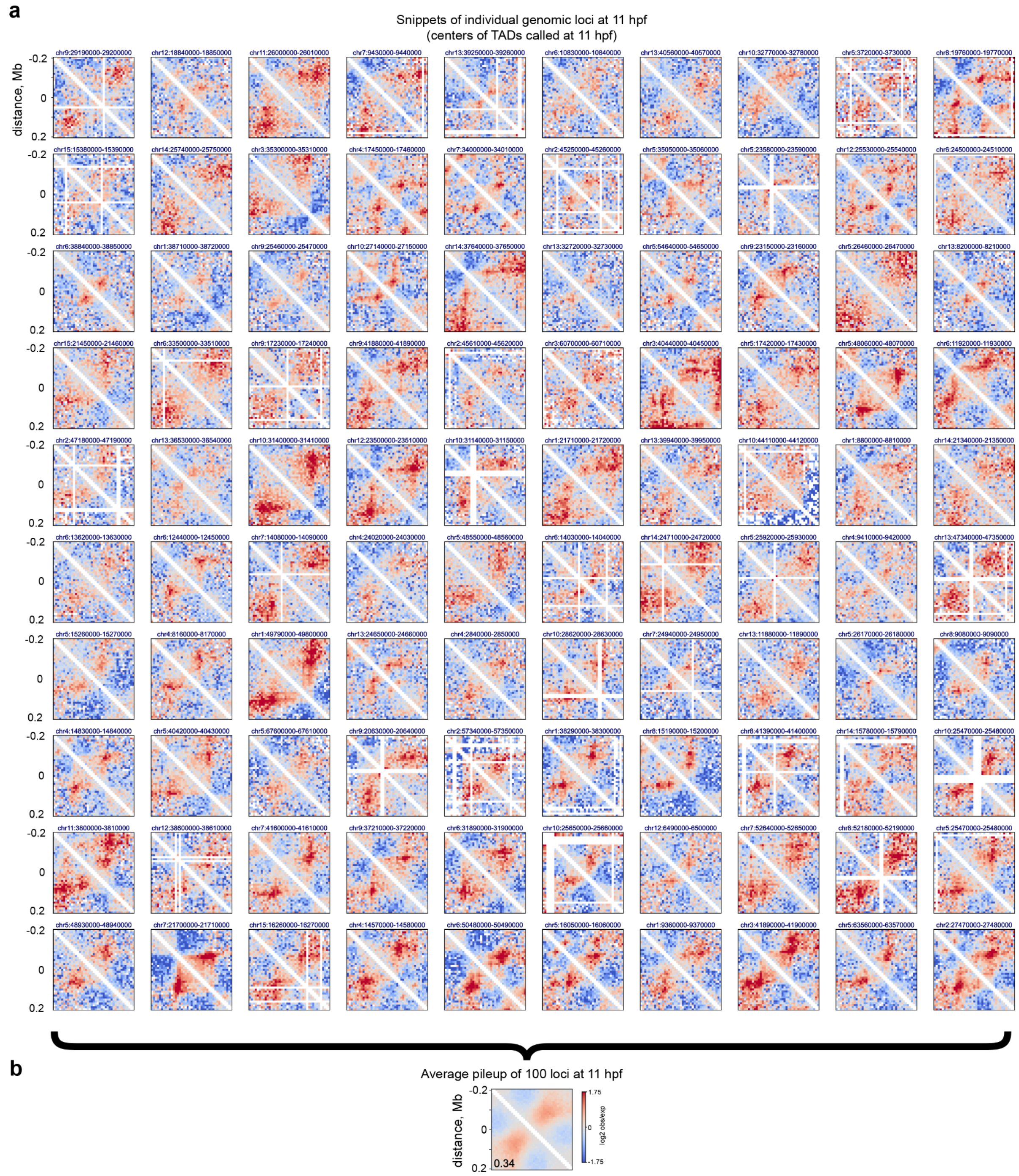

**Figure SI 2. Averaging centers of TADs at 11 hpf.**

a. Snippets of Hi-C maps of 100 centers of TADs called at 11 hpf. Each snippet is named by the genomic position of the TAD center.

b. Average pileup with fountain-like-looking output. The average does not resemble individual conformations. The number in the corner represents the strength of the fountain signature.

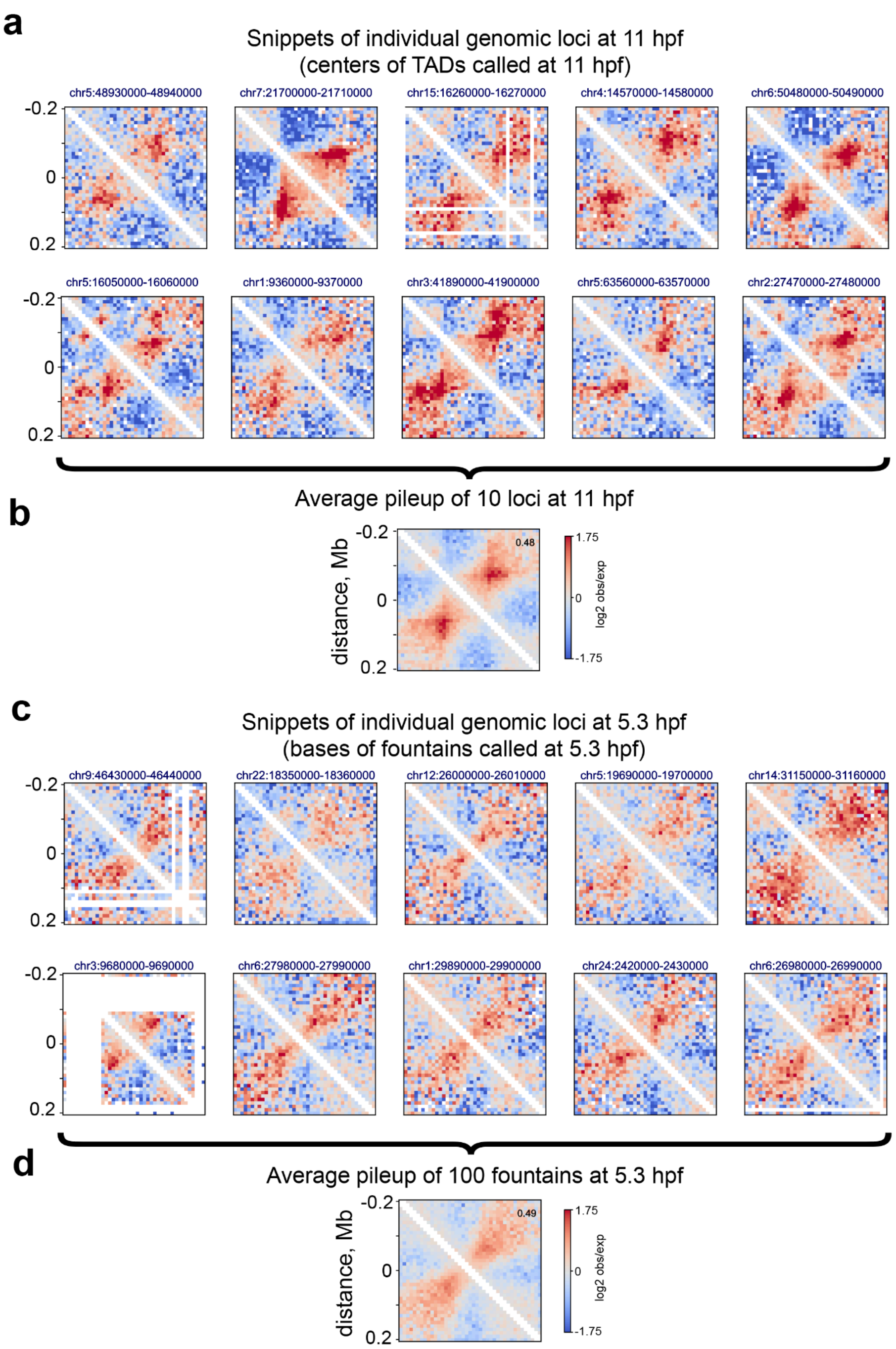

**Figure SI 3. Principles of dealing with averaging error in Hi-C: visual inspection of the snippets, averaging over the snippets with features of similar sizes.**

a. Snippets of 11 hpf Hi-C maps of 10 centers of TADs called at 11 hpf, all TADs have similar sizes. Averaging results in average TAD.

b. Snippets of 5.3 hpf Hi-C maps of 10 fountain bases called at 5.3 hpf. Averaging results in average fountain.

#### III. Validations and characteristics of fountains

##### III.1. Genomic view with two fountains at 5.3 hpf of zebrafish development

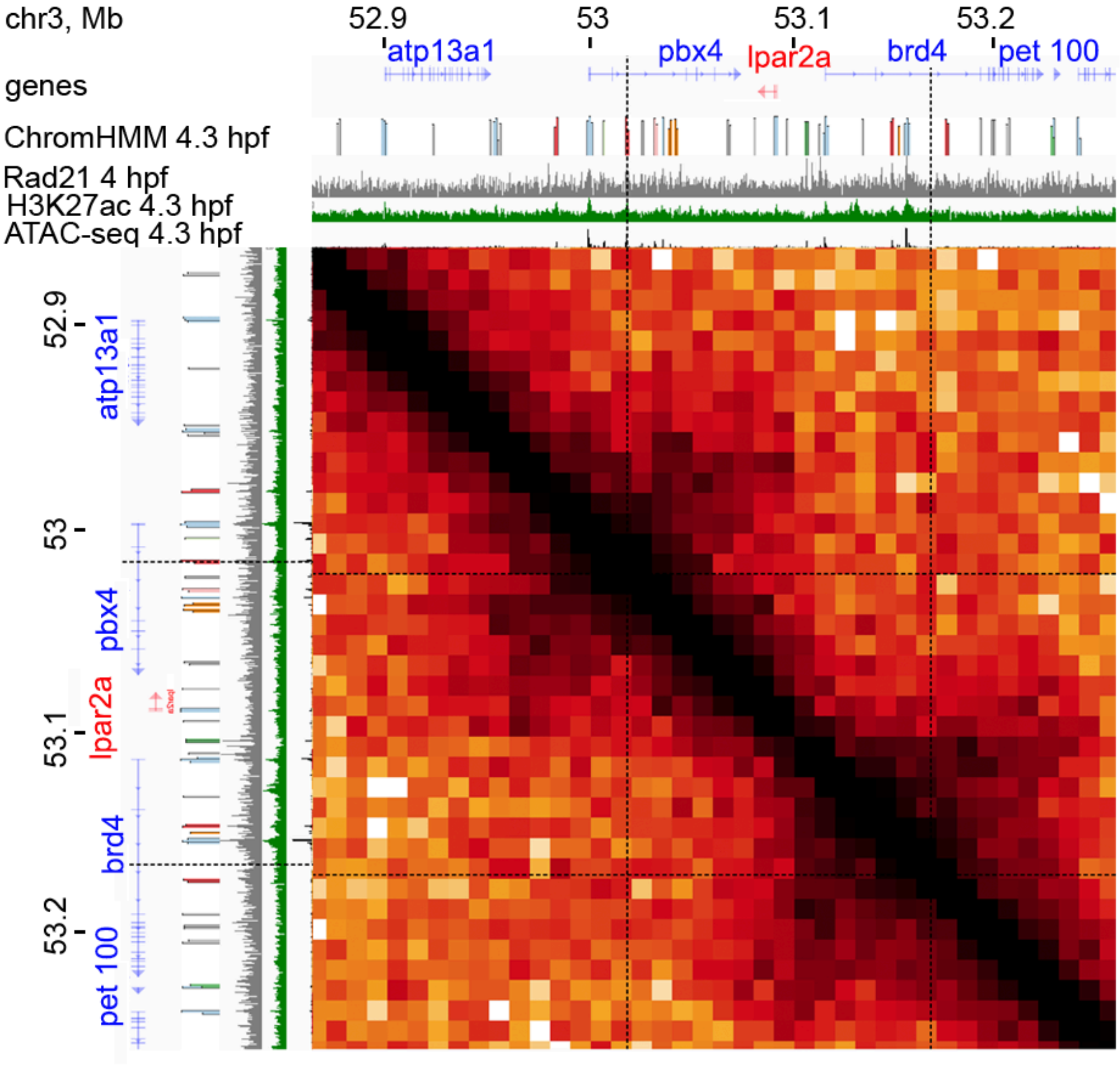

**Figure SI 4.** HiGlass [24] genome browser view on two fountains with gene track, annotation of developmental regulatory elements (ChromHMM at dome ATAC-Seq peaks from [25]), and epigenetic annotations. Hi-C bin size is 10 Kb. The colors of regulatory elements correspond to the Main Fig. 2c (TSS A1 and A2 are blue, TSS Flank 1 and 2 are green, Enhancer is red, Enhancer flank is rose, Enhancer Weak 1 is orange, Poised is purple, Repressed is light-purple, and Quiescent is gray).

#### III.2. Fountains are associated with zygotic transcription

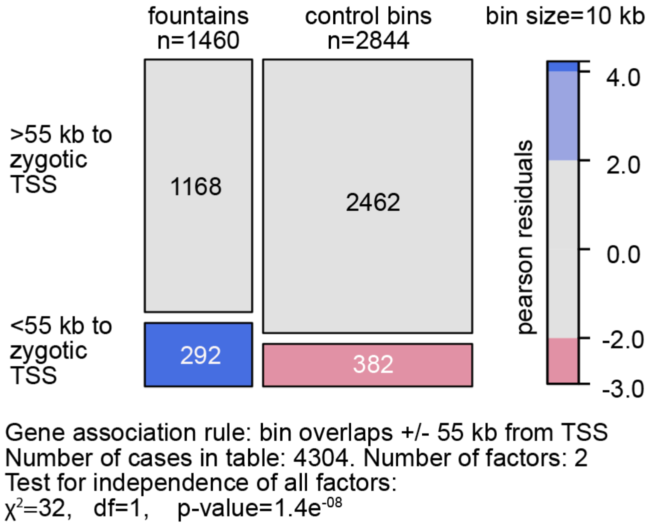

**Figure SI 5. Fountains preferentially localize near zygotic genes.**
Chi-squared test. Control 10 kb bins were taken at the distance +/- 1 Mb from 1460 fountains, the overlapping fountain bins were excluded. The number of zygotic genes in the embryo (4777) and RNA-seq data are from [26].

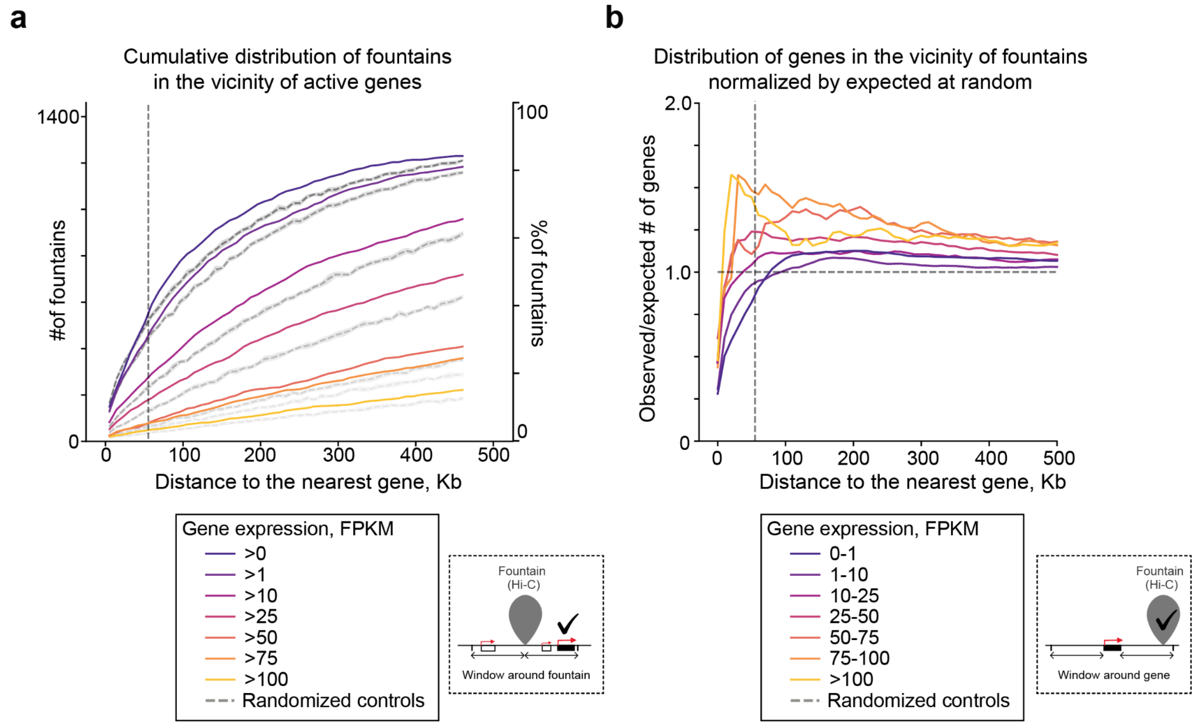

**Figure SI 6. Fountains preferentially localize near active zygotic genes and vice versa: active genes localize near fountains.**
**a**. Cumulative distribution of fountains in the vicinity of active genes. Gene expression is from EBI expression atlas [27]. Potential maternal transcripts were excluded from the analysis

(maternal transcripts from [26]). Control: randomly picked genomic regions. Note that genes of expression level category are enriched over control (100 randomizations of fountain positions).
**b**. Distribution of genes near fountains normalized by expected at random. Horizontal line: line of no enrichment. Above the horizontal line: enrichment over expected; below the horizontal line: depleted. Vertical line: 55 Kb distance line as in SI Fig. 5 above.
Note that more actively transcribed genes are enriched in fountains at distances below 100 Kb near them. Genes with little or no expression tend to have fewer fountains around them at distances below 50-100 Kb than expected at random.

#### III.3. Protractor tool

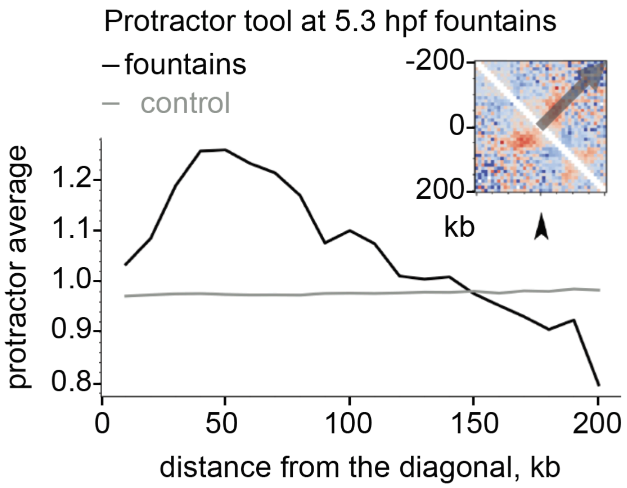

**Figure SI 7. Application of protractor tool [11] for validation of fountains.**
The grey arrow in the inset (fountain) represents the perpendicular to the main diagonal (ideal hairpin).
The protractor shows the mean values in Hi-C snippets at fountains along the represented direction. The protractor average has a clear hump at 50 Kb, which means that the high observed over expected number of interactions at fountains, perpendicular to the main diagonal.

The grey line is a control based on non-fountain genomic regions, which means the absence of increase of interactions at short ranges in random genomic bins.

#### III.4. Fountains are independent of TAD centers

With Nikolai Bykov

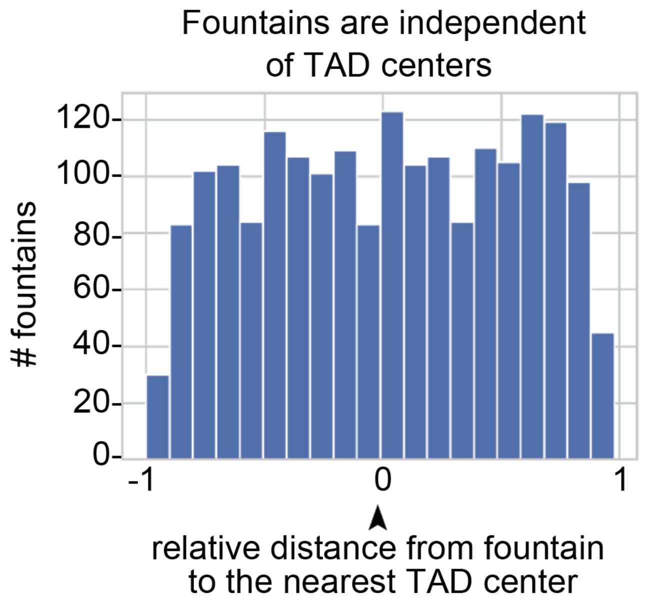

**Figure SI 8. Distribution of fountains around centers of TADs: no correlation was found.**

To determine individual TAD, we took all the boundaries at 11 hpf, confirmed by CTCF presence at this stage, and paired sequential pairs of boundaries. We then compared the positioning of the fountains to the TAD centers and plotted the distributions of distance between fountains to the nearest TAD center.

### IV. Detailed characterization of epigenetics at zebrafish fountains

With Kristina Perevoshchikova

To characterize the epigenetics at fountains in zebrafish embryogenesis, we first studied the distribution of chromatin openness and epigenetics factors at fountain bases around 4-5 hpf (Fig SI 4a). As a control, we compared these distributions to non-fountain 10 Kb bins. We ranked the factors by the significance of the difference in means between fountains and non-fountains. Fountains have high H3K27ac, PolII binding, chromatin openness, Nanog and Sox2 binding, Rad21 enrichment, H3K4me1, Pou5f3, and p300. H3K4me3 is not significantly different at fountains than in other genomic locations. CTCF is significantly depleted at fountains.
Next, we asked what factors are associated with the change of the fountain strength in the MZ*triple* (Fig SI 4b). For that, we calculated the deltas of fountain scores between wild-type cells and MZ*triple* (as the differences between values). We correlated them with the deltas for epigenetic factors available from [28]. We found that ATAC-Seq, H3K27ac, and H3K4me1 changes have the highest correlation with the fountain strength change, while H3K4me3 has correlation close to 0.

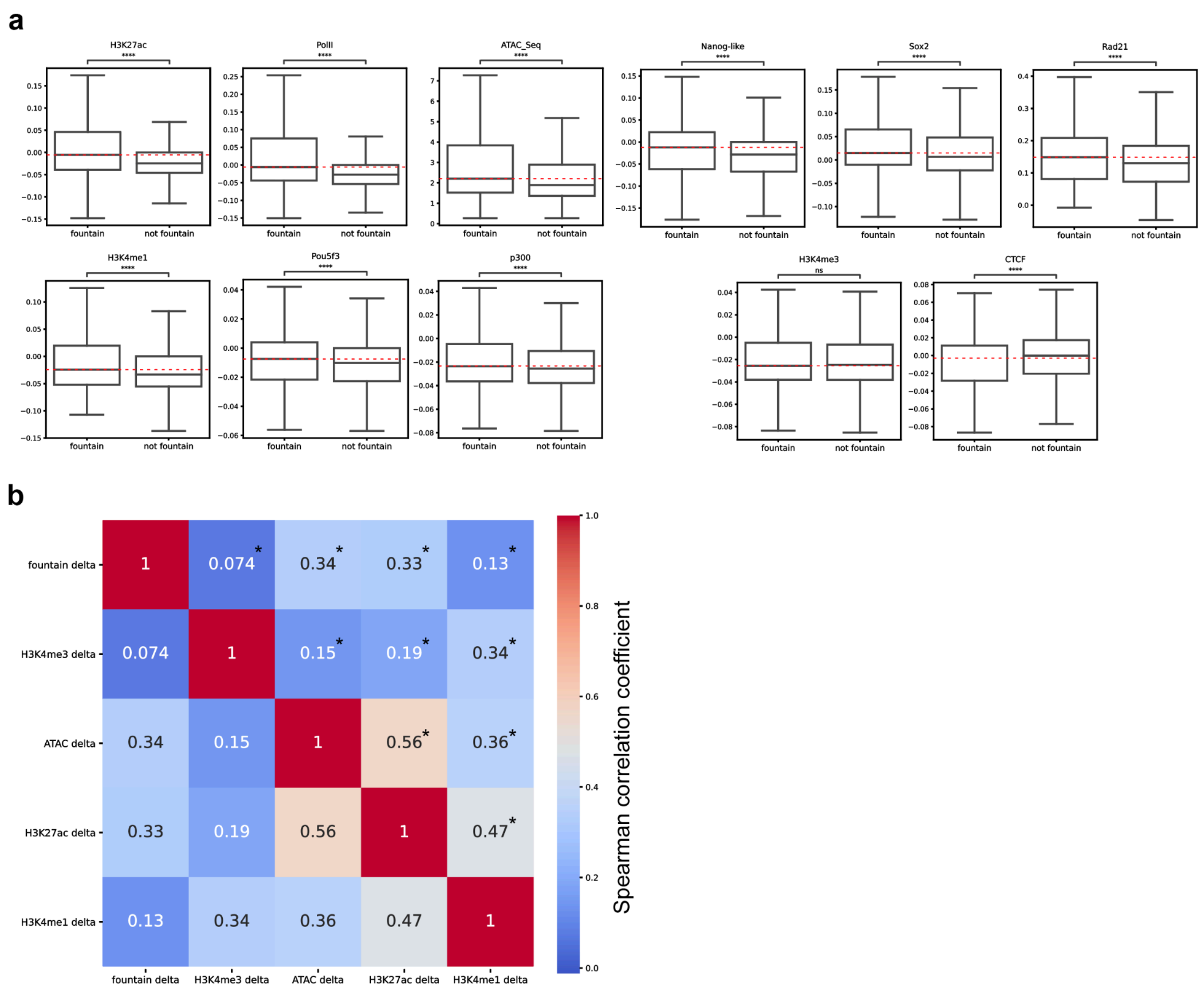

**Figure SI 9. Factors associated with fountains and fountain strength**

**a.** (Related to Figure 3a) Boxplots of epigenetic tracks at fountains versus non-fountain bins with similar levels of chromatin openness, ordered by the significance of the difference in means (from

left upper corner to right bottom corner). Only those characteristics that are positively associated with fountains are shown (p-value<0.05 for the Mann-Whitney test with a greater alternative after Bendjamini-Yakuteli multiple testing correction). The bins next to the fountains were removed from the analysis. The last two boxplots represent H3K4me1, which is not significantly associated with fountains, and CTCF, which is negatively associated with fountains. The red dashed line shows the mean distribution for fountain bins. The datasets and hours are the same as in Figure 3a and Ext. Data Fig. 2a-c.
**b.** (Related to Figure 6) Spearman correlation of epigenetic changes with fountain score change in MZ*triple* mutant. *- P-value of Beta-test implemented in *scipy* < 0.01.

#### V. Fountain calling in medaka and *Xenopus*

For fountain calling in other species, we first re-mapped the data from Niu et al., 2021 [29] (*Xenopus tropicalis*, genome xenTro10) and Nakamura et al., 2021 [8] (*Oryzias latipes*, medaka fish, genome oryLat2) with Open2C *bwa-mem* [30] and *pairtools*[31]-based pipeline *distiller-nextflow* version 0.3.3 and walks policy “all”.
With the visual inspection of Hi-C maps, we confirmed the presence of fountain patterns at 5 kb resolution for medaka at developmental stage 10 and 10 kb resolution for *Xenopus* at developmental stage 11. We filtered out poorly mapped genomic regions at these resolutions (as was done for zebrafish) and ran *fontanka* with the reference fountain mask from zebrafish.
We then filtered out fountain peaks that were too weak and had large noise scores (filters iv.a, iv.b, iv.f from “Fontanka protocol” section of Methods).
Dot patterns and genomic misassemblies more frequently contaminated the resulting fountains in *Xenopus* and medaka (Fig SI 5-6) than those in zebrafish (Fig SI 1). This can be attributed to (1) worse quality of the genome assemblies for these species than for zebrafish; and (2) the presence of dots and TADs at selected developmental stages alongside the fountains.
Although the average fountain, based on the calls, displayed a prominent fountain signature (Figure 9), it is important to confirm that individual snippets also resemble fountains (see section I. “Emergence of a fountain as an average pileup” of this Supplemental Information).

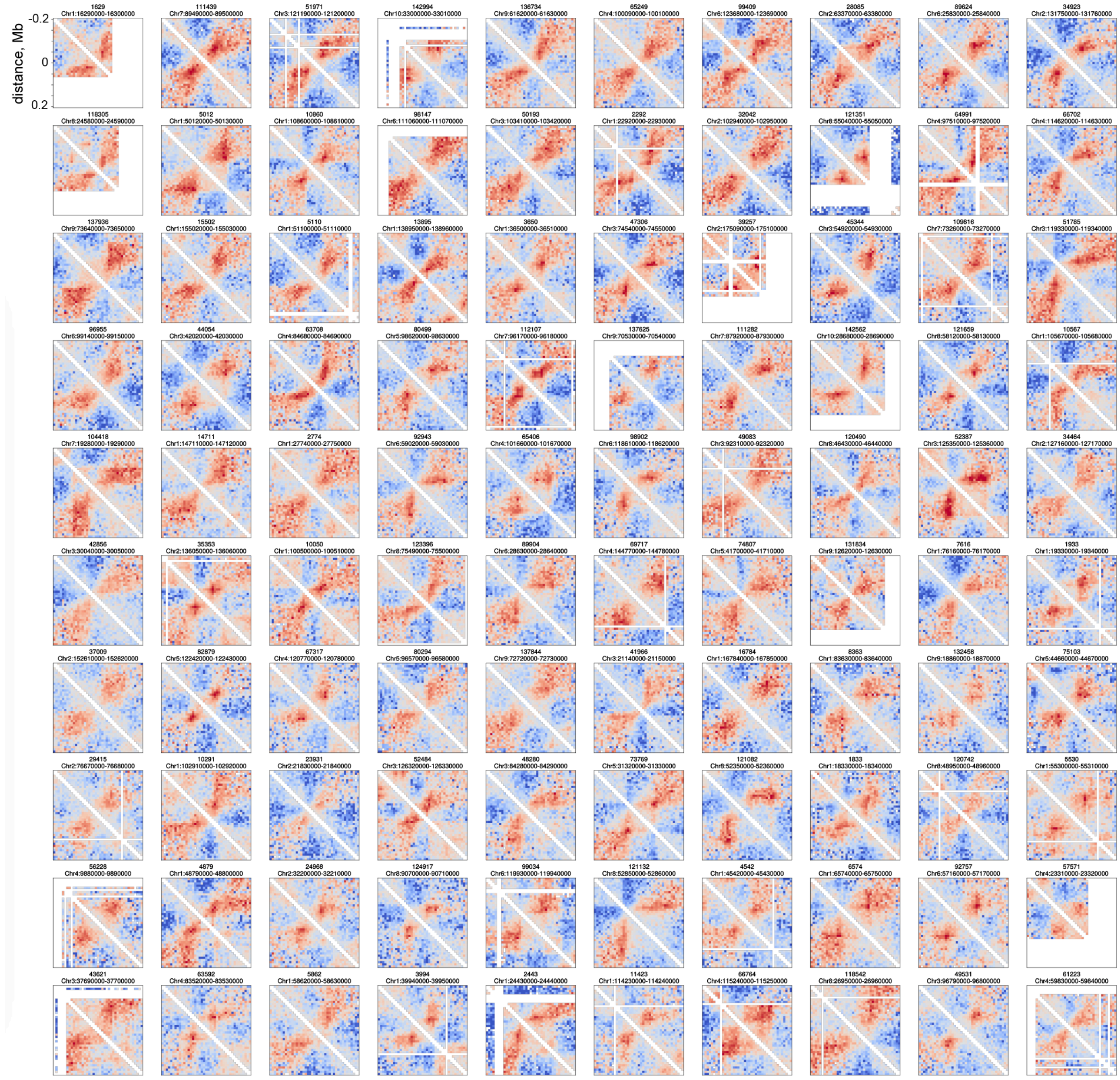

**Figure SI 10. Individual fountains of *Xenopus tropicalis* at developmental stage 11.**
Snippets of Hi-C maps of 100 fountain bases (top by fountain score with the reference fountain) called with *fontanka*. Each snippet is named by the genomic bin index of the fountain base and the genomic position of the fountain base.

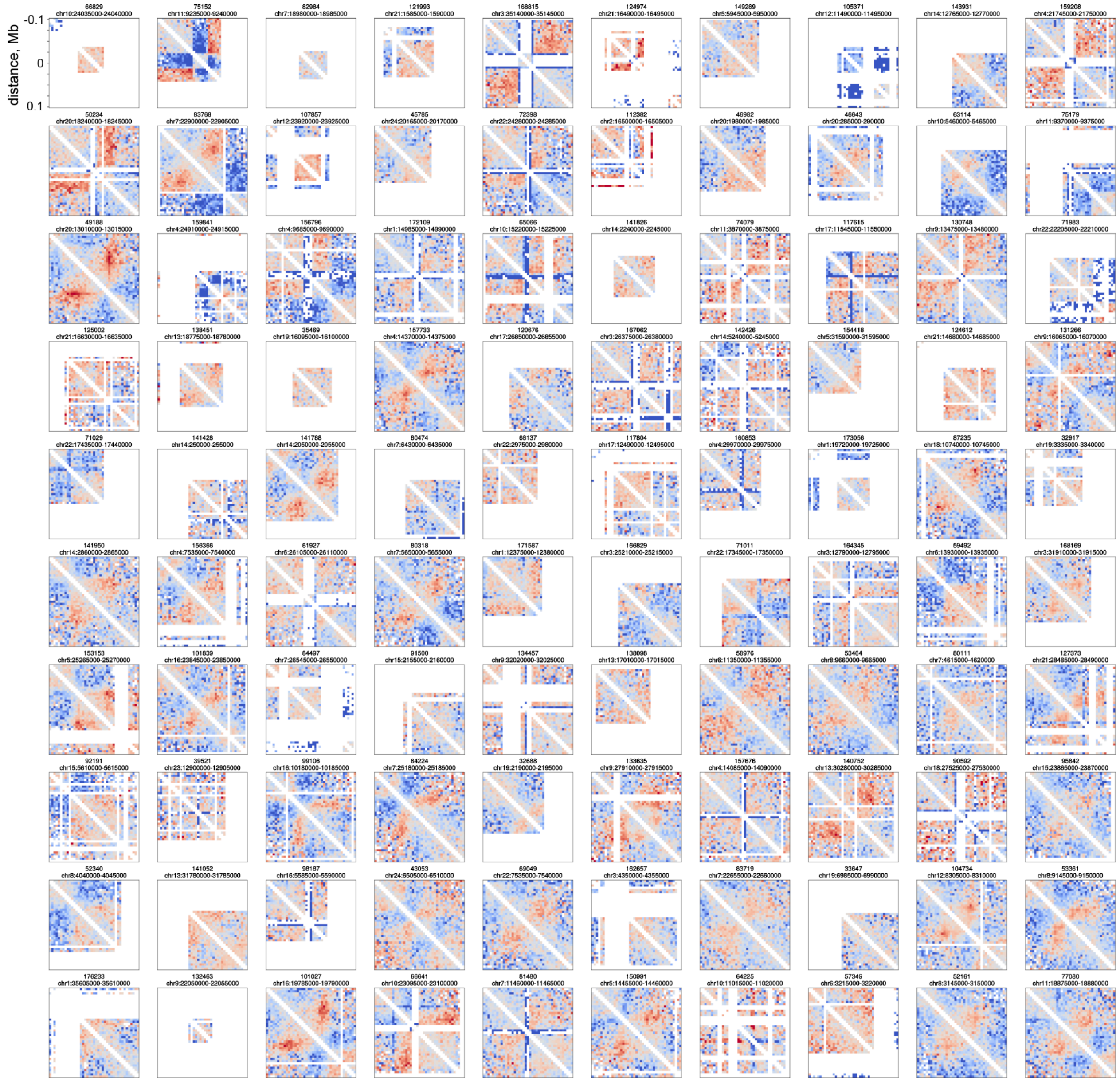

**Figure SI 11. Individual fountains of medaka fish at developmental stage 10.**
Snippets of Hi-C maps of 100 fountain bases (top by fountain score with the reference fountain) called with *fontanka*. Each snippet is named by the genomic bin index of the fountain base and the genomic position of the fountain base. Note that true fountain calls are often contaminated with dots and poorly assembled regions.

### VI. Individual fountains in mouse cell cycle

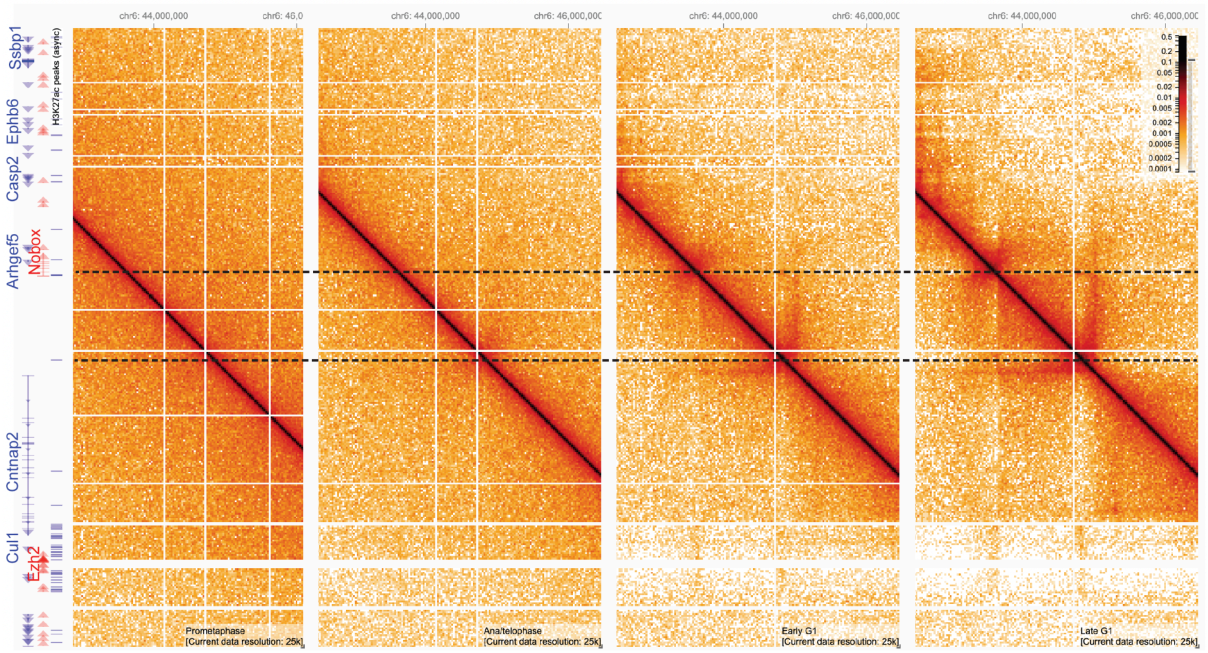

**Figure SI 12. HiGlass [24] genome browser view on the genomic region with two fountains in mouse G1E erythroblast cell line emerging in the cell cycle.**

(*Central*) Hi-C for four stages of cell cycle from [32] with (*left*) gene track and H3K27ac peaks from [33]. Mouse genome coordinates (mm10) are shown above the maps. Hi-C bin size is 25Kb. Dashed lines mark the locations of two fountains emerging at early G1 and later transforming into other chromatin structures (TAD and stripe, correspondingly). Note that fountain 2 demonstrates a fountain signature at the ana/telophase already.
